# Quasi-universal scaling in mouse-brain neuronal activity stems from edge-of-instability critical dynamics

**DOI:** 10.1101/2021.11.23.469734

**Authors:** Guillermo B. Morales, Serena Di Santo, Miguel A. Muñoz

## Abstract

The brain is in a state of perpetual reverberant neural activity, even in the absence of specific tasks or stimuli. Shedding light on the origin and functional significance of such a dynamical state is essential to understanding how the brain transmits, processes, and stores information. An inspiring, albeit controversial, conjecture proposes that some statistical characteristics of empirically observed neuronal activity can be understood by assuming that brain networks operate in a dynamical regime near the edge of a phase transition. Moreover, the resulting critical behavior, with its concomitant scale invariance, is assumed to carry crucial functional advantages. Here, we present a data-driven analysis based on simultaneous high-throughput recordings of the activity of thousands of individual neurons in various regions of the mouse brain. To analyze these data, we synergistically combine cutting-edge methods for the study of brain activity (such as a phenomenological renormalization group approach and techniques that infer the general dynamical state of a neural population), while designing complementary tools. This strategy allows us to uncover strong signatures of scale invariance that is ”quasi-universal” across brain regions and reveal that all these areas operate, to a greater or lesser extent, near the edge of instability. Furthermore, this framework allows us to distinguish between quasi-universal background activity and non-universal input-related activity. Taken together, this study provides strong evidence that brain networks actually operate in a critical regime which, among other functional advantages, provides them with a scale-invariant substrate of activity covariances that can sustain optimal input representations.

## 1. Introduction

The brain of mammals is in a state of continuous ongoing activity even in the absence of stimuli or specific tasks [1, 2, 3]. Shedding light onto the origin and functional meaning of such an energy-demanding baseline state of background dynamics and its interplay with input-evoked activity are challenging goals, essential to ultimately understand how different brain regions represent, process and transmit information [4, 5, 6]. Following the fast-paced development of powerful neuroimaging and electrophysiological technologies such as two-photon calcium imaging and neuropixels probes, recent years have witnessed important advances in our understanding of these issues.

An inspiring hypothesis —which aims to become a general principle of brain dynamical organization— posits that neuronal networks could achieve crucial functional advantages, including optimal information processing and transmission, by operating in the vicinity of a critical point [7, 8, 9, 10, 11]. Using the jargon of statistical physics, this implies that the network operates in an intermediate regime at the border between ”ordered” and ”disordered” phases [7, 10, 12, 13, 14, 15, 16, 17, 18, 19, 20, 21]. Criticality, with its concomitant power-laws, entails well-known functional advantages for information processing such as, e.g., an exquisite sensitivity to perturbations and a huge dynamic range [12, 11] to name a few. Moreover, it also gives rise to the emergence of a broad spectrum of spatio-temporal scales, i.e. scale invariance or simply ”scaling” [11, 22, 23], that could be essential to sustain brain activity at a broad hierarchy of scales, as required to embed highly-diverse cognitive processes.

In spite of its conceptual appeal and thrilling implications, the validity of this so-called ”criticality hypothesis” as an overarching principle of dynamical brain organization remains controversial [24, 25, 26]. Therefore, novel theoretical approaches and more-stringent experimental tests are much needed to either prove or disprove this conjecture and, more in general, to advance our understanding of the mechanisms underlying brain ongoing activity.

From the theoretical side it is crucial to refine the conjecture itself and discern what type of criticality is the most pertinent to describe brain activity [8, 11]. Different possible scenarios have been explored; among them [11]: (i) the edge of activity propagation, with scale-free avalanching behavior [7, 10, 14, 27], (ii) the edge of a synchronization phase transition [28, 21, 29, 30, 31, 11], and (iii) the edge of a stable-unstable transition, often called ”edge of chaos” but to which we will refer as ”edge of instability” hereon [32, 33, 34].

From the empirical side, evidence of putative brain criticality often relies on the detection of scale-free bursts of activity, called ”neuronal avalanches” [7, 35, 17, 36, 29, 37], the presence of robust long-range spatio-temporal correlations [38, 39], the analysis of large-scale whole-brain models fitted to match empirically observed correlations [40, 41, 42, 43], etc. Recently, with the advent of modern techniques enabling simultaneous recordings of thousands of neurons, complementary experimental evidence revealing the existence of scaling in brain activity —which might or might not stem from underlying criticality [8, 11, 26, 44]— has emerged from unexpected angles. Among these, let us mention (i) a novel renormalization-group approach which identified strong signatures of scale-invariant activity in recordings of more than a thousand neurons in the mouse hippocampus [45, 46], (ii) the direct inference —using linear response theory and methods from the physics of disordered systems— of ”edge-of-instability” type of critical behavior in neural recordings from the macaque monkey motor cortex [34], and (iii) the discovery of an unexpected power-law distribution, which revealed scale invariance in the spectrum of the covariance-matrix from tens of thousands of neurons in the mouse visual cortex while the mouse was exposed to a large set of sequentially-presented natural images [6].

What all these works have in common, is the fact the they use specifically-designed, powerful mathematical tools to analyze vast amounts of high-throughput data.

Here we bring these diverse approaches to a common ground and develop novel and complementary tools leveraging them synergistically in order to analyze state-of-the-art neural recordings in diverse areas of the mouse brain [47]. As shown in what follows, these analyses strongly enhance our understanding of scale-invariance and possible criticality in the brain. In particular, they allow us to elucidate the emergence of quasi-universal scaling across regions in the mouse brain, to conclude that all such regions are, to a greater or lesser extent, posed close to the edge of instability, as well as to disentangle background and input-evoked neural activities.

### Theoretical framework and open-ended questions

For the sake of self-containedness, let us briefly discuss the three innovative theoretical approaches cited above, along with some stemming open-ended questions that we pose (further technical details are deferred to the Methods section).

#### (A) Renormalization-group approach to neuronal activity

The renormalization group (RG) is retained as one of the most powerful ideas in theoretical physics, allowing us to rationalize collective behavior —at broadly-diverse observational scales— from the properties of the underlying ”microscopic” components, and to understand, for instance, the emergence of scale invariance [23]. In a remarkable contribution, Meshulam *e*t al. developed a phenomenological RG approach to analyze time series from large populations of simultaneously recorded individual spiking neurons and scrutinize their collective behavior [45, 46]. The method, similar in spirit to Kadanoff’s blocks for spin systems [23], allows one to construct effective descriptions of time-dependent neural activity at progressively larger ”coarse-grained” scales. Notably, a number of non-trivial features —generally attributed to scale-invariant critical systems— emerge from the application of such RG analyses to recordings of more than 1000 neurons in the mouse hippocampus while it is moving in a virtual-reality environment. These features include among others: (i) a non-Gaussian (fixed-point) distribution of neural activity at large scales, (ii) non-trivial scaling of the activity variance and auto-correlation time as a function of the coarse-graining scale, and (iii) a power-law decay of the spectrum of the covariance matrix [45, 46, 48].

A limitation of this type of phenomenological RG analysis is that, even if it is capable of uncovering scale invariance, it does not allow discerning what kind of putative phase transition could be at its origin. Moreover, doubts have been raised about the possible interpretation of the results as stemming from criticality (see [49, 50] and below).

In any case, leaving aside for the time being these caveats, one can wonder whether the observed scaling features are shared by other brain regions or if they are rather specific to the mouse hippocampus. Is there any kind of ”universality or at least ”quasi-universality” (in the sense of scaling exponents showing limited variability) in the neural dynamics across brain regions despite their considerable anatomical and functional differences? Is it possible to find empirical evidence that allows us to discern whether the observed scaling actually stems from critical behavior?

### (B) Inferring the dynamical regime from neural recordings

Dahmen *et al*. devised a general approach —based on linear-response theory ideas and tools from the physics of disordered systems— that allows one to infer the overall dynamical state of an empirically observed neuronal population. In particular, this theoretical approach permits us to estimate the distance to the ”edge of instability” from empirical measurements of the mean and dispersion of ”spike-count covariances” (also called ”long-time-window” covariances) across pairs of recorded neurons [34]. Straightforward application of this approach to neural recordings from motor cortex of awake macaque monkeys at rest strongly supports the idea that such a region operates in a ”dynamically balanced critical regime with nearly unstable dynamics” [34]. Thus, one can wonder whether other brain regions —for instance, the hippocampus in (A)— can also be empirically proven to be similarly close to the edge of instability.

Moreover, recently Hu and Sompolinsky went a step further and derived an analytical expression (based also on linear-response theory for large random networks) for the full spectrum of eigenvalues of the spike-count covariance matrix as a function of its distance to the edge of instability. This provides us with an alternative method to estimate the distance to the edge of instability from empirical data. In particular, these analyses reveal that the eigenvalue distribution develops a power-law tail as the network approaches the edge of instability [51]. The resulting non-trivial eigenvalue distribution stems from the recurrent network dynamics near the edge of instability [51] and it clearly differs from the Marchenko-Pastur law for the correlations of independent random units [52]. Thus, we also infer what is the distance to the edge-of-instability in the theoretical model that best fit the spectra of spike-count covariance matrices obtained from empirical data.

### (C) Scaling in optimal input representations

Stringer *et al*. [6] studied experimentally and theoretically the spectrum of the covariance matrix in neuronal populations in mouse primary visual cortex (VISp) while the mouse was exposed to a very large set of sequentially-presented natural images (recording more than 10^4^ neurons in parallel; see Methods). From the resulting data they found that the spectrum of the covariance matrix obeyed ”*an unexpected power law* ” [6]: the n-th rank-ordered eigenvalue scaled as 1*/n*^*µ*^ with *µ ≳* 1.

This power-law decay of the rank-ordered eigenspectrum was somehow surprising; the authors were expecting a much faster decay, as would correspond to a lower-dimensional representation of the visual inputs (see below). Notice that, here, by ”representations” one means neural activity that stems from or is correlated with sensory or task-related inputs. On the other hand, the term ”dimensionality” is employed in the sense of ”principal component analysis” (PCA) [53], where the dimension is the number of principal components required to explain a given percentage of the total variance; often most of the variability in neural data can be recapitulated in just a few principal components or dimensions [54], but this is not always the case [6].

Remarkably, Stringer *et al*. were also able to prove that the power-law decay of the rank-ordered eigenvalues is not an artifact directly inherited from the statistics of the inputs, but a very deep mathematical property: it stems from a trade-off between the neural representation of visual inputs being as high-dimensional as possible (i.e. including a large number of non-negligible components in a PCA analysis) and mathematically preserving its smoothness (i.e. its continuity and differentiability). As a simple illustration of this last abstract property, let us mention that the smoothness of the representation prevents, for instance, that tiny variations in the inputs dramatically alter the neural population activity, which translates into a more robust encoding [6].

Let us finally recall that common knowledge in statistical physics tells us that a power-law decay of the covariance-matrix spectrum (i.e., of the ”*propagator* ”) is one of the most remarkable generic trademarks of critical behavior, emphasizing the emergence of a scale-free hierarchical organization of spatio-temporal correlations [23]. Indeed, this spectrum is one of the objects studied in the phenomenological RG approach (A), revealing a power-law decay (for the mouse hippocampus) with an exponent *µ <* 1 [45], which violates Stringer *et al*.’s bound for continuous neural representations (*µ ≳* 1).

The results by Stringer *et al*. trigger a cascade of questions: is the empirically-observed scaling of the spectrum of the covariance matrix a mere consequence of the external input being represented in an optimal way? In other words, does the covariance spectrum obey scaling also in the absence of inputs, i.e. for resting-state or ongoing activity? Is intrinsic criticality in the network dynamics required to ”excite” such a broad spectrum of modes supporting optimal input representations? How come the exponent *µ* measured by Meshulam et al. in the hippocampus is smaller than 1 in seeming contradiction with Stringer *et al*.’s predictions?

Summing up: while (A) allows us to detect and scrutinize the presence of scaling in empirical recordings in a systematic and quantitative way, (B) provides us with practical tools to infer the actual dynamical regime of the underlying neural network, thus paving the way to ascribe empirically-reported scaling to edge-of-instability criticality, and (C) sets the scene for relating criticality to so far unexplored functional advantages for optimal input representation.

In what follows, we use these methods in a synergetic way while developing novel and complementary tools to scrutinize scale invariance and criticality across regions in the mouse brain, thus providing data-driven answers to most of the previously raised questions.

## Results

Most of the forthcoming analyses rely on the empirical electrophysiological data presented by Steinmetz *et al*. in [47], where the activity, *x*(*t*), of thousands of individuals neurons (in particular, the precise times of their spikes) is simultaneously recorded at a high 200Hz resolution in several mouse brain regions (as illustrated in Fig.1). These recordings include periods in which the mouse is performing some specific task and some in which is at a ”resting-state”. Therefore, we first separate both types of time series and analyze the corresponding ”resting-state” and ”task-related” activity independently, paying special attention to the former. In addition, we also consider data from recordings of mouse VISp from Stringer *e*t al. [6] (see Methods). In all cases, we restrict our analyses to areas (see Fig.1) with at least *N* = 128 simultaneously-recorded neurons.

**Figure 1:**
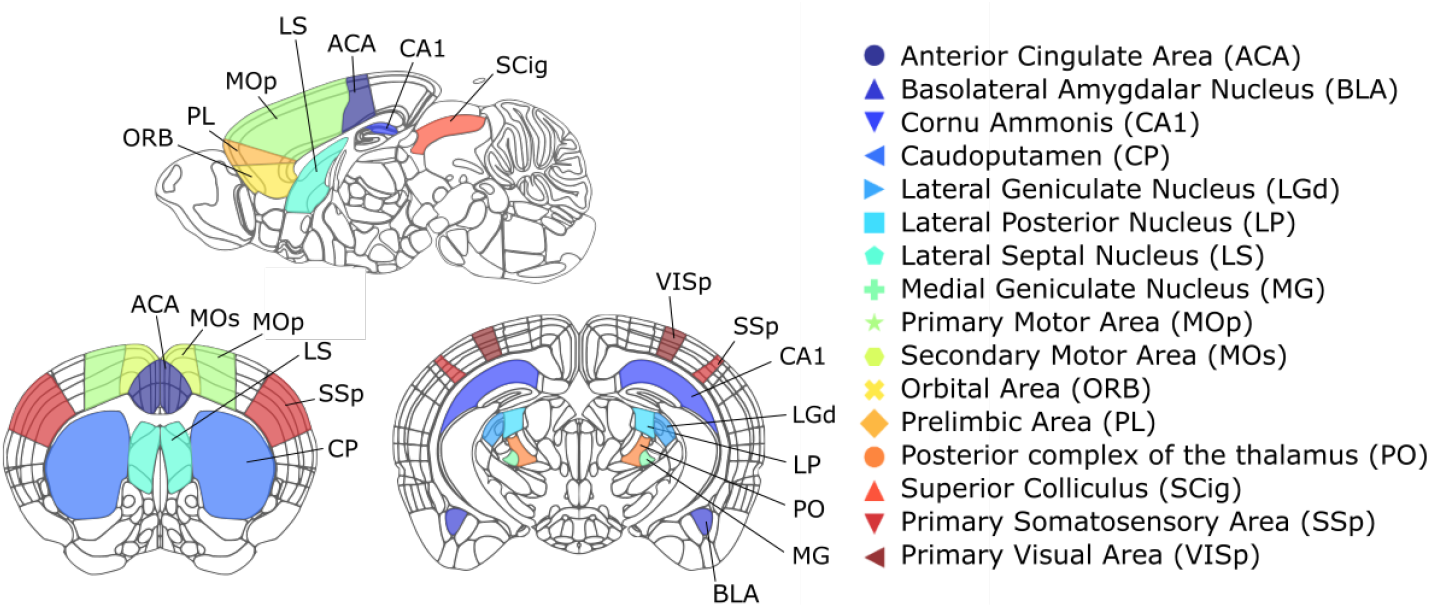
Schematic representation of the regions in the mouse brain —using three different projections— considered in this work, together with their names and corresponding acronyms. Credit: Allen Institute, Atlas brain maps: https://atlas.brain-map.org/atlas.

### Quasi-universal scaling across brain regions

We first employ the phenomenological RG approach to scrutinize whether non-trivial scaling behavior, such as the one reported in [45, 46] for the CA1 region of mouse hippocampus, is observed in other areas of the mouse brain. For the sake of clarity let us summarize the gist of the RG approach (further details in the Methods Section as well as in [45, 46]), along with our main results.

Following the spirit of Kadanoff’s block one seeks to perform a coarse graining of *N* ”microscopic variables” —i.e. single-neuron activities in this case— to construct effective descriptions at progressively larger scales. Nevertheless, given the absence of detailed information about physical connections (synapses) between neurons, a criterion of maximal pairwise correlation (rather than the standard one of maximal proximity) is used to block together pairs of neurons in a sequential way [45]. Let us remark that, for this, it is crucial to determine pairwise correlations in a careful and consistent way; this requires the choice of a suitable discrete time bin for each data set, for which we have devised an improved protocol (see *Time-scale determination* in the Extended Methods of the SI). In this way, the activity time series of the two most correlated neurons are added together and properly normalized, giving rise to effective time series for ”*block-neurons*” or simply ”*clusters* ” of size 2. One then proceeds with the second most-correlated pair of neurons and so on, until all neurons have been grouped in pairs. The process is then iterated in a recursive way, so that after *k* coarse-graining (RG) steps there remain only *N*_*k*_ = *N/*2^*k*^ block-neurons, each recapitulating the activity of *K* = 2^*k*^ individual neurons.

The distributions of activity values across block-neurons at level *k, P*_*k*_({*x*}), can be then directly computed for the different steps of the RG clustering procedure and, from them, a number of nontrivial features can be identified for all the considered mouse brain areas [47]:

#### Non-Gaussian probability distribution of block-neuron activity

Fig.2A shows the probability distribution *Q*_*k*_(*x*) of non-zero activity values, *x*, of the coarse-grained block-neurons at one of the RG steps (*k* = 5, i.e., *K* = 32). Observe also, in the inset of Fig.2A, the curve collapse obtained for sufficiently large values of *K* as well as the presence of significant non-Gaussian tails, as exemplified for one of the considered brain regions: the primary motor cortex (MOp). This convergence of *Q*_*k*_(*x*) to an asymptotic shape after a few RG steps, as observed for all regions (see Fig.S1 in the SI), suggests that a nontrivial fixed point of the RG flow has been reached and that the emerging distribution of activity is rather universal across brain regions in resting conditions.

**Figure 2:**
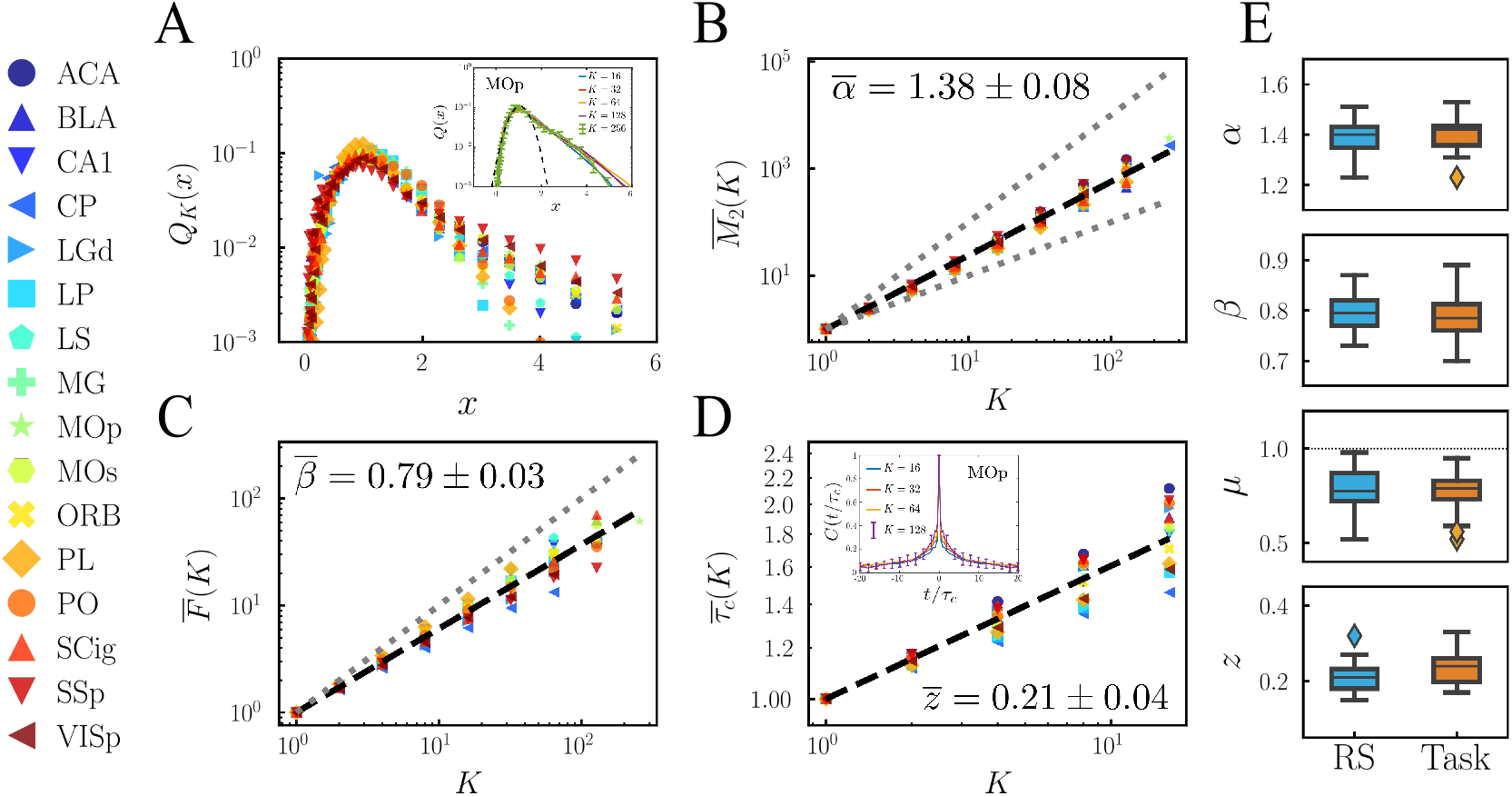
Results of the phenomenological RG analyses of brain activity measured in 16 different mouse brain areas (A-D resting state activity). (**A**) Probability distribution for normalized non-zero activity in block-neurons of size *K* = 32 across brain regions (main panel), as well as (inset) at 5 consecutive steps of the coarse-graining for a representative region (MOp). (**B**) Variance of the non-normalized activity as a function of the block-neuron size, *K*, in double logarithmic scale (upper and bottom dotted lines, with slopes 2 and 1, mark the fully correlated and independent limit cases, respectively). (**C**) Scaling of the free energy *F*_*k*_ as defined in Methods (the dotted line corresponds to the expected behavior for un-correlated variables). (**D**) Inset: decay of the autocorrelation function as a function of the rescaled time *t/τ*_*c*_ for the MOp region; a different value of *τ*_*c*_ is used for each cluster size but, after rescaling, all data collapse into a common curve. Main plot: scaling of the characteristic correlation time *τ*_*c*_ as a function of *K* in double logarithmic scale for the different areas. To facilitate the comparison between areas with different number of neurons, the variance has been normalized as 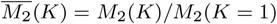, the probability of being silent re-scaled to 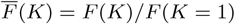 and the correlation time as 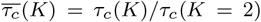. (**E**) Comparison of the exponent values *α, β, µ* and *z* for resting-state (RS) and task-related activity (Task). Horizontal line inside each box represents the sample median across regions, the whiskers reach the non-outlier maximum and minimum values, and the distance between the top (upper quartile) and bottom (lower quartile) edges of each box is the inter-quartile range (IQR). Outliers are represented with a diamond marker. For the exponent *µ*, the critical value *µ*^*c*^ = 1 has been marked with a dotted line.

#### Scaling of the activity variance

As shown in Fig.2B an almost perfect scaling is observed for the variance *M*_2_(*K*) of the non-normalized activity of block neurons, with an average exponent 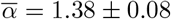 across regions. In particular, we measure *α*_*CA*1_ = 1.37 ± 0.03 ± 0.02 (mean + mean-absolute-error of individual measurements + standard deviation over experiments, see Methods) for the CA1 region within the hippocampus, which is within errorbars of the value *α* = 1.56 ± 0.07 ± 0.16 reported in [46] for the same area. Notice that all these exponent values are always in between the expected ones for uncorrelated (*α* = 1) and fully-correlated variables (*α* = 2), revealing consistently the existence of non-trivial scale-invariant correlations.

#### Scaling of the ”free-energy”

This is defined as *F* (*K*) = − log(*S*_*K*_), where *S*_*K*_ is the probability for a block-neuron of size *K* to be silent within a time bin. As shown in Fig.2C, this quantity exhibits a clear scaling with cluster size, with an average exponent 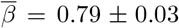 across regions (0.78 ± 0.04 ± 0.05 for CA1, to be compared with the value 0.87 ± 0.014 ± 0.015 reported in [46]).

#### Scaling of the auto-correlation time

Fig.2D shows that, despite the very broad variability of intrinsic timescales for individual neurons within each region (see Fig.S3 in the SI), dynamical scaling can be observed in the decay of the block-neuron autocorrelation times *τ*_*c*_(*K*) (see Methods) in all regions, with an average exponent across regions 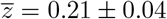 (with *z*_*CA*1_ = 0.18 ± 0.03 ± 0.01 for CA1, in perfect agreement with the one reported in [46] for this region (*z* = 0.22 ± 0.08 ± 0.10) and compatible also with the exponent values reported in [44] using a different approach). As an additional test for dynamical scaling, we show in the SI (Fig.S2) how the curves for the auto-correlation functions at different coarse-graining levels collapse when time is appropriately re-scaled.

#### Scaling of the covariance-matrix spectrum

By diagonalizing the covariance matrix computed at different levels of coarse-graining, it is possible to analyze how their corresponding spectra decay with the rank of the eigenvalues and how their cut-offs change with cluster size. As illustrated in Fig.3, in all the analyzed brain areas there is a clear power-law scaling of the eigenvalues with the rank, with an average exponent across regions 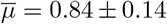, as well as a common dependence with the fractional rank (*rank/K*), the latter manifested in the collapse of the curves at different levels of coarse graining, much as in [45]. Likewise, the value reported in [45] for CA1 (*µ* = 0.76±0.05±0.06) is in perfect agreement with our measured value for the same region, *µ*_*CA*1_ = 0.78±0.08±0.02. Although we will address this point later on, let us for now stress that *µ* is smaller or, at most, approximately equal to one 1 in all regions, seemingly suggesting discontinuity of the neural representations [6].

**Figure 3:**
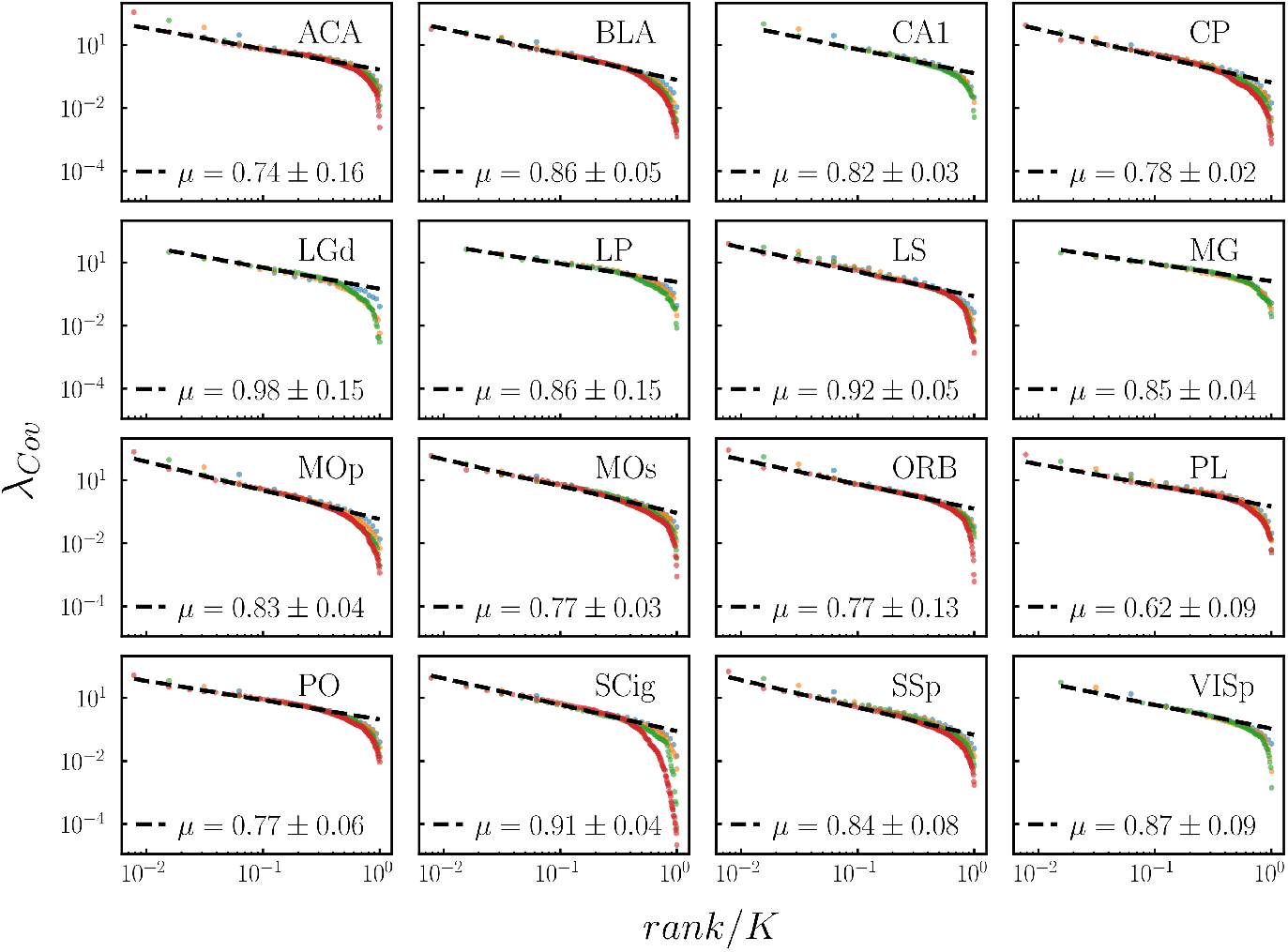
Scaling of the covariance matrix spectrum for the resting-state activity in clusters of size *K* = {16, 32, 64, 128} (blue, yellow, green and red markers, respectively) in 16 different brain regions. Observe not only the decay of the rank-ordered eigenvalues as a power law of the rank, but also the excellent collapse of the cut-offs obtained after rescaling the eigenvalue rank by the total size *K*. For each region, we collect the average powerlaw exponent and its standard deviation across experiments (see also Table S2 in the SI).

For the sake of consistency, we have also verified that the reported exponent values exhibit little variability upon changes in the time-discretization bin, with the exception of the exponent *µ*, for which we observe an increase on longer time-scales beyond the typical inter-spike-interval of the population activity (see Fig.S5 in SI).

Moreover, as it turns out, similar signatures of scale-invariance to those reported for resting-state activity emerge in RG analyses of neural recordings obtained while the mice are performing a task (see section *Datasets* in Extended Methods of the SI and [47] for more details). This similarity is illustrated in Fig.2E, which shows how the dispersion and mean value of the scaling exponents across regions are not significantly altered (*p >* 0.1 on a two-sample t-test for each exponent) when one compares the resting-state and task-related activity (see also Fig.S6).

Finally, as a control test we verified that the non-trivial scaling features revealed by the RG analyses are lost for all areas when the correlation structure of the data is broken either by: (i) reshuffling the times of individual spikes in the time series; (ii) shifting each individual time series by a random time span while keeping the sequence of spikes; or (iii) shuffling spikes across neurons (see Figs.S12-S14 in SI).

### Dynamical state of resting-state activity across brain regions

Despite the elegance and appeal of the RG results —as originally presented by Meshulam *e*t al.— their straightforward interpretation as stemming from underlying criticality has been questioned [49, 50]. In particular, very similar scaling behavior was found to emerge in a (non-critical) model of uncoupled neurons exposed to latent correlated inputs [49] (see Discussion). Thus, it is not guaranteed *a priori* that the scaling behavior we just found across brain regions stems from underlying criticality and the RG approach does not allow us to provide an answer to this question. Therefore, we resort to alternative methods to estimate the dynamical regime of each brain region from empirical data as described above [34, 51]. For this, one needs to compute spike-count covariance values across pairs of neurons, which measure the pairwise correlations in time-integrated activity (i.e. the total number of spikes) across samples (see Methods).

In particular, to compute meaningful covariances one assumes that neural activity is stationary, a condition that typically holds during recordings of resting-state type of activity, but not as much in task-related activity, for which network activity is inherently input-driven and non-stationary. Therefore, we restrict our forthcoming analyses to resting-state data. We also notice that the values of spike-count covariances depend on the window sizes over which such counts are measured [34]. In the forthcoming results, we took sampling windows of *T* = 1*s*, which are sufficient for autocorrelations to decay while maximizing the number of samples over which spike-count covariances are computed (see Fig.4A an Table S1 in the SI). Using such pairwise spike-count covariances measured in a given region, we consider two alternative methods to infer the distance of its dynamical state to the edge of instability:

**Figure 4:**
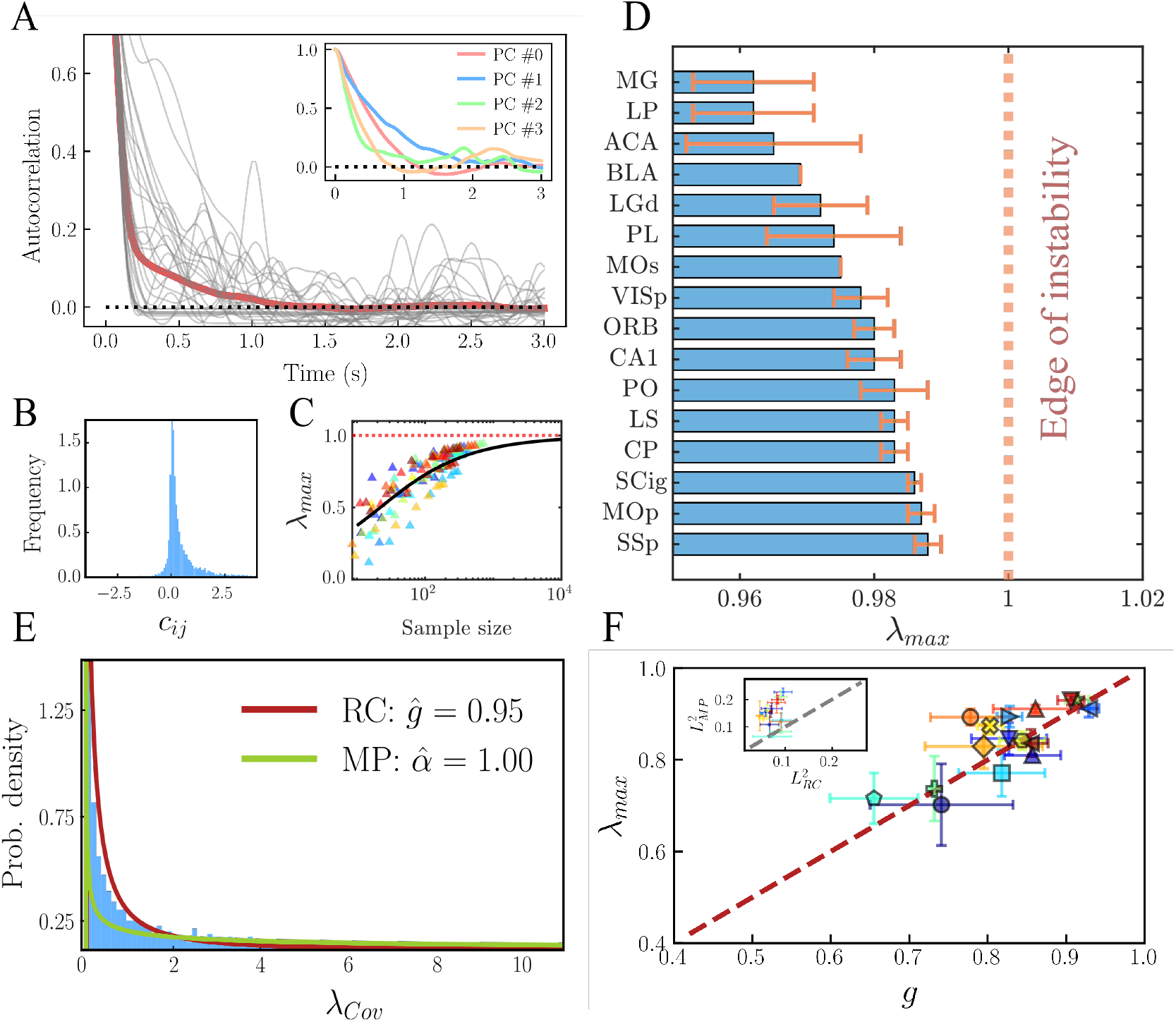
(**A**) Autocorrelations decay in the MOp region (grey lines correspond to 30 randomly chosen neurons, while red line denotes the average across all neurons). Notice how autocorrelations vanish on a timescale of ∼ 1*s*. Inset: Decay of the average autocorrelation for the firing rates projected into the first four principal components. (**B**) Distribution of pairwise covariance values in one of the considered brain regions (MOp). Observe the strong peak around 0 (as expected from theoretical approaches of ”balanced networks” [55]) as well as the large dispersion of values [34, 51], including (asymmetric) broad tails. (**C**) Dependence of the estimated value of *λ*_*max*_ on the number of neurons *N* in the sample. Each color corresponds to a region following the color code in Fig.1; for each region we show 10 points, corresponding to sub-samples of 10% to 100% of the total neurons recorded, using the experiment with a greater number of recorded neurons for each area. Fitting the resulting values of *λ*_*max*_ for different values of *N* to Eq.1 with a parameter Δ common to all regions and extrapolating for large values of *N*, one observes a fast convergence to the very edge of instability. (**D**) Proximity to criticality in 16 different regions of the mouse brain, as measured by the estimated largest eigenvalue *λ*_*max*_ —with *λ*_*max*_ = 1 marking the ”edge of instability”— of the inferred connectivity matrix. The regions are ordered according to their average *λ*_*max*_ and the error bars are calculated as standard deviation over different experiments in the same region. (**E**) Long time-window covariance eigenvalues distribution for an example region (MOp, blue histogram), together with the best-fitting Marchenko-Pastur distribution (green line, fitted parameter 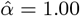 = 1.00) and the best-fitting covariance eigenvalues distribution for a linear-rate model of randomly connected neurons (red line, fitted parameter 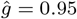 = 0.95). (**F**) Fitted values of *g* and *λ*_*max*_ (including errorbars, calculated as in D) for all the considered regions. Inset: deviation of the empirical distribution to the Marchenko-Pastur (MP) distribution versus deviation to the theoretical distribution for a recurrent network (RC) of linear rate neurons close to the edge of instability.

#### (i) Estimations from the statistics of spike-count-covariance distributions

Calling the mean value of the distribution of such spike-count covariances 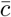 and its standard deviation *δc*, the largest eigenvalue of the connectivity matrix, *λ*_*max*_ —whose closeness to 1 marks the distance to the edge of instability, thus gauging the dynamical regime— can be determined from the following expression [34]:

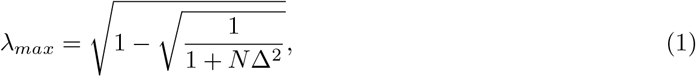

where *N* is the total number of neurons in the population, 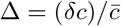. We remark that the derivation of Eq.(1) is based on linear response theory and relies on tools from spin-glass theory applied to networks of spiking neurons, which assumes the stationarity of the spiking statistics [34]. An Augmented Dickey-Fuller test was performed to verify in each case the validity of this assumption (see Table S1 and section *Stationarity of spiking statistics* in Extended Methods of the SI).

As illustrated in Fig.4B (for one of the selected regions), the distributions of pairwise spike-count covariances turn out to be sharply-peaked around 0, but they exhibit significant tails, revealing the presence of heterogeneously correlated pairs in all areas.

From these spike-count covariances, one could then easily estimate the distance to the critical point for the neural dynamics in each region (see Table S1 and Fig.4F). Let us remark, however, as it follows from Eq.1, that the value of *λ*_*max*_ strongly depends on the number of neurons being recorded and, since the available empirical data heavily sub-samples each region, the aforementioned approach would actually underestimate the real value of *λ*_*max*_, which becomes much closer to 1 in the limit of tens of thousands of neurons. In Fig.4C we plot the values of *λ*_*max*_ as a function of the number of recorded cells, showing that the larger the neural population size, the larger the value *λ*_*max*_.

Moreover, because of this dependence with the system size, it becomes difficult to compare the estimated values across regions with different numbers of neurons recorded. To avoid this limitation, we computed for each experiment and region the normalized covariance width, Δ*t*, and then applied Eq.1, extrapolating to a common number of neurons *N* = 10^4^. In Fig.4D we show the distance to the critical point thus estimated, with errorbars computed as the standard deviations across experiments in each region. Notice that most values lie on a very narrow window between 0.96 and 0.99, with a mean value 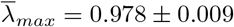 = 0.978 ± 0.009, close to the edge of instability.

#### (ii) Estimations from the eigenvalue spectrum

Hu and Sompolinsky [51] have recently derived an analytical expression for the distribution of eigenvalues of the spike-count covariance matrix for a recurrent random network of linear rate neurons, whose dynamics follows the equation:

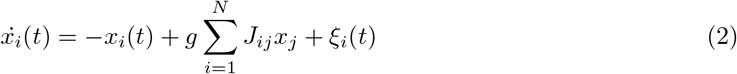

with ⟨*ξ*_*i*_(*t*)*ξ*_*j*_(*t* +*τ*_0_)⟩ = *σ*^2^*δ*_*ij*_*δ*(*τ*_0_) and *J*_*ij*_ ∼ 𝒩 (0, 1*/N*). In the model, the parameter *g* sets the overall connection strength, with 1 − *g* marking the distance to the edge of instability [51]. This result allows one to estimate the network dynamical state by fitting the actual empirically-determined spectrum to the theoretical distribution as a function of *g* (see Extended Methods in the SI and [51]). In particular, Fig.4E shows the best fit (*g* = 0.95) of the empirically-determined eigenvalue distribution for the MOp region to the Hu-Sompolinsky distribution, together with a (much worse) fit to the Marchenko-Pastur distribution (expected for random uncorrelated time series), thus underlying the non-trivial structure of the observed spike-count covariances, which stem from recurrent quasi-critical interactions. A summary of the inferred *g*-values for all areas is shown in Fig.4F (see also Table S1), which further illustrates the strong similarity between the results obtained with the two employed methods, when the original number of neurons in each experiment is considered.

### Non-trivial scaling features emerge in the RG analysis of recurrent random-network models at the edge of instability

To close the loop, we wondered whether a simple model of randomly-coupled linear units, as the one defined by Eq.(2), is able to reproduce also other non-trivial scaling features as revealed by phenomenological-RG analyses. In Fig.S10 of the SI we show evidence that such recurrent neural networks driven by noise generate patterns of activity that reproduce many of the non-trivial scaling features emerging out of the RG analyses when they are posed close to the edge of instability (e.g., *g* = 0.95), while such non-trivial features are lost in the subcritical regime (where trivial Gaussian scaling emerges). In particular, the values of the exponents at the edge of instability (*α* ≈ 1.28, *z* ≈ 0.21 and *µ* ≈ 0.78) are remarkably similar to the ones measured across regions in the mouse brain (see Table S2). The observation is rather striking given that these exponent values come from a linear model of randomly connected neurons, while real data are, most likely, generated by a more complex non-linear dynamics on heterogeneous networks. A recent work on universal aspects of brain dynamics [56] might help shedding light onto this result.

### Non-universal, input-dependent exponents in task-related activity

Saying that external stimuli shape neural correlations in information processing areas is, to a certain extent, an obvious statement. However, the fact that the spectrum of the activity covariance matrix follows a very simple mathematical rule that ultimately ensures the smoothness of the internal representation of inputs —as mathematically proved by Stringer *et al*. [6]— is quite remarkable. In particular, these authors showed that the neural representation for a *d*-dimensional (visual) input —that is to be encoded collectively in the neural activity of the primary visual cortex (VISp)— is constrained by the requirement of smoothness of the representation (i.e., continuity and differentiability of the associated representation manifold) [6]. Being more specific, for these conditions to hold, the spectrum of the covariance matrix needs to decay as a power-law with an exponent *µ* that must be greater than 1 for continuity, and greater than 1+2*/d* —where *d* is the dimension of the input ensemble (see *Datasets* section in Extended Methods of the SI)— for differentiability [6]. Thus, for sufficiently ”complex” inputs, i.e. with large embedding dimensionality (*d*), this exponent is constrained to take a value arbitrarily close to (but larger than) unity, *µ ≳* 1.

On the other hand, in the previous RG analyses, we found values of *µ* consistently smaller than 1 —both under resting conditions and in task-related activity— for all the considered areas (see Fig.3), in agreement with [45], but in seeming contradiction with the predictions of Stringer *et al*. [6]. How come that we report values *µ <* 1 in all areas including sensory information encoding ones, even when the mouse is exposed to external stimuli? Does it mean that stimuli encoding violates the requirements for efficient representation put forward by Stringer *et al*. [6]?

At the core of this seeming paradox lies a data-processing method proposed by Stringer *e*t al. that allows one to extract the input-only related covariances from the overall ”raw” covariance matrix [6]. The approach stems from the idea that population activity can be decomposed into an input-related (or ”input-encoding”) subspace, which spans input-only related activity and a complementary space —orthogonal to the former one— which captures the remaining activity [57]. In a nutshell, the so-called cross-validated PCA (cv-PCA) method consists in repeating twice the very same experiment and comparing the two resulting raw covariance matrices; this comparison allows one to infer which part of the covariance is shared by the two matrices and is, hence, input-related (see secion 6 in the SI) and which complementary part stems from unrelated background activity [6]. Unfortunately, given the nature of such an experimental protocol, it is not possible to apply cv-PCA to the dataset of Steinmetz *e*t al. [47] extensively employed above.

However, to prove that our RG results above (consistent with *µ <* 1) are not in contradiction with the ones by Stringer *e*t al. (*µ* ≥ 1), we have extended the cv-PCA method (as explained in detail in section 6 of the SI) to be able to actually extract from empirical data in [6] not just the input-related covariance matrix but also the time-series of input-related neural activity. For this, the overall activity *x*(*t*) of a given neuron at time *t* is projected into two separate sub-spaces, i.e. decomposed as:

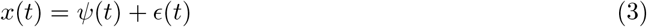

where *ψ*(*t*) describes its input-related activity and *E*(*t*) stands for the remaining “orthogonal” activity, responsible for trial-to-trial variability. This decomposition allows us to perform separate RG analyses to input-related and background activity data. As a consistency check, we also verified that the covariance matrix eigenspectrum associated with *ψ*(*t*) has no significant difference with the one obtained from the standard application of cv-PCA analyses in [6] (see Fig.S15 in the SI for further details).

We now proceed to show the main results of applying the phenomenological RG approach over the data of [6] considering: *(i)* the raw timeseries *x*(*t*), *(ii)* the input-related activity *ψ*(*t*), and *(iii)* the background activity *E*(*t*), with the results averaged over three different mice.

i. Analyzing the raw data (i.e. *x* variables) one observes again exponent values (*α* = 1.49 ± 0.08 and *µ* = 0.73 ± 0.08) in agreement with the previously-reported quasi-universal values for different mouse-brain areas (see Fig.5, together with Fig.S7 and Table S1 in the SI), highlighting the robustness of our results for very different recording techniques.
ii. Considering the input-evoked activity (*ψ*) one finds that the exponents *α* —which was rather robust when measured from the raw-activity data— is significantly altered (see SI, Fig.S7). More importantly, a significant increase in the exponent *µ* is observed with respect to the raw-data analyses; it now respects the theoretical boundary *µ >* 1 + 2*/d* (with *d* → ∞ in natural images) for the smoothness of the representation manifold (see Fig.5). In addition, the value of *µ* obtained from our RG analyses decreases as the input dimensionality grows, in agreement with the theoretical results and empirical findings in [6].
iii. Finally, considering only the residual ”orthogonal” parts (*ϵ*) we found slightly smaller values of *µ* with respect to the raw data analyses for the two types of inputs considered (see Fig.5). We notice, however, that this background activity is not to be confused with simple ”white noise” as it exhibits correlations with a non-trivial power-law spectrum. When looking at the exponent *α* for the variance inside block-neurons, we did not observe a significant change with respect to the overall-activity case (Fig.S7 in SI). This further supports the idea that the observed scaling exponents in the overall raw data are dominated by the higher-dimensional, background activity.

**Figure 5:**
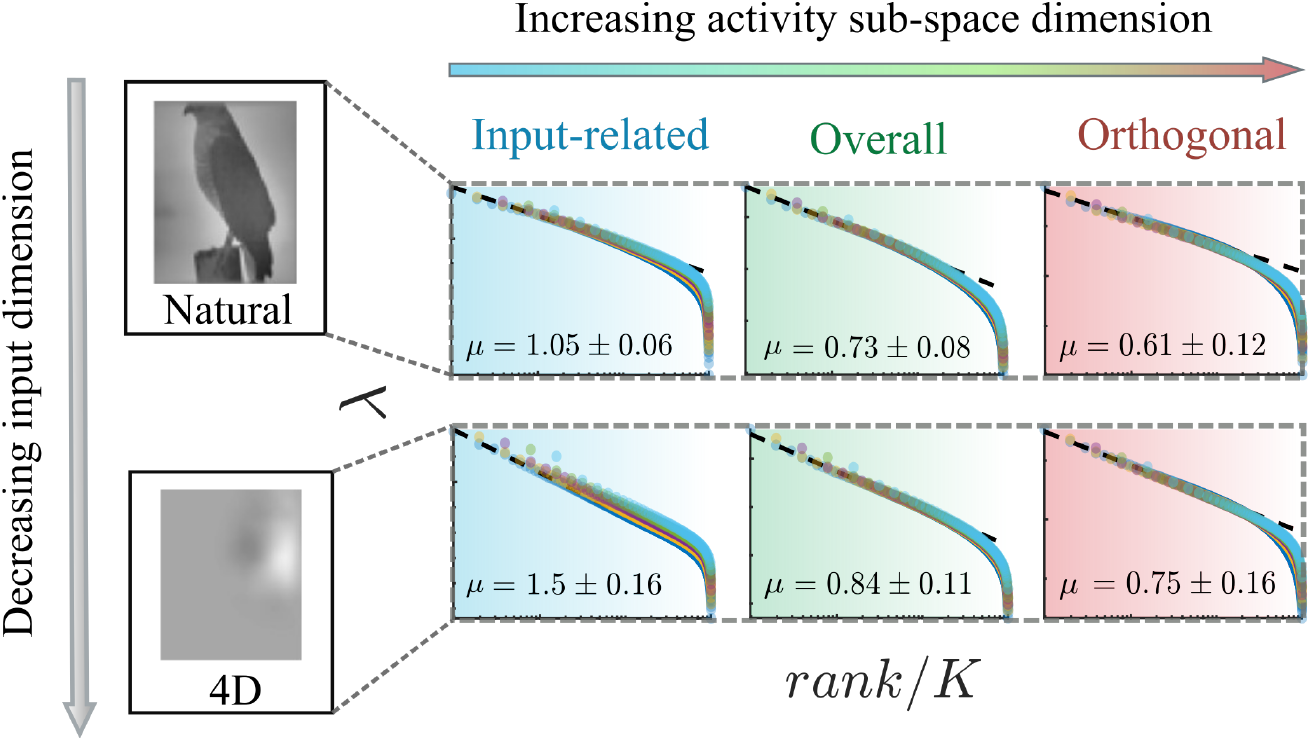
Diagram showing the two observed trends in the power-law exponent *µ*, which characterizes the decay of the covariance-matrix eigenvalues as a function of their rank. *µ* decreases with the complexity of the input when activity is projected into the task-encoding subspace (also called representation manifold). On the other hand, for a fixed type of input, the exponent decreases with the proportion of background, ”noisy” activity in the data, which lies in a higher-dimensional subspace orthogonal to the representational manifold. Natural and low-dimensional images examples have been adapted from [6].

To summarize the previous results, Fig.5 recaps the two observed trends in the scaling exponent *µ*. On the one hand —for data obtained using images of different intrinsic dimensionalities (see Extended Methods in SI)— we found the relationship predicted and observed by Stringer *et al*. in [6], namely, that for the input-related activity *ψ*, the value of *µ* decreases with the complexity of the input signal, with a lower limit of *µ* = 1 for high-dimensional inputs. On the other hand, the exponent *µ* decreases with the relative weight of background activity in the data: the more input-related activity that is projected away, the flatter the spectrum of the remaining orthogonal space.

As a side note, let us mention that —owing to the nature of the experimental setup in [6], in which stimuli are interspersed with grey-screen inter-stimulus intervals — computation of actual dynamical correlations are not meaningful in this case, so that we purposely left aside the study of the dynamical scaling exponent *z*. Likewise, the transformations carried on the overall data to extract the input-related and orthogonal activity do not necessarily preserve the biological significance of the zero-activity, making the computation of the ”free-energy” exponent *β* pointless.

Thus, in summary, application of the phenomenological RG procedure to the data for VISp of Stringer *e*t al. [6] reveals that the overall activity exhibits clear signatures of scale invariance, sharing its quasi-universality with the previously analyzed regions, which relied on a different dataset. Nevertheless, this overall activity can be decomposed into input-related and noisy/orthogonal activities: the scaling exponent for the covariance matrix in the first case obeys the mathematical constraints derived by Stringer *e*t al., thus solving the seeming contradiction with the spectrum of the overall data.

## Discussion

Understanding how the brain copes with inputs from changing external environments, and how information from such inputs is transmitted, integrated, processed, and stored in a physical substrate consisting of noisy neurons —exposed also to a stream of other overlapping inputs— is one of the major challenges in neuroscience. The fast development of revolutionary experimental and observational techniques as well as of novel theoretical insights have resulted in huge advances in this field in recent years.

On the one hand, novel technologies allowing for the simultaneous recording of thousands of neurons, pave the way to quantitative analyses of brain activity with unprecedented levels of resolution and detail, making it possible, for the first time, to discriminate between different overarching theories.

Here, we have taken advantage of high-throughput data, together with state-of-the-art theoretical approaches to analyze neuronal activity across regions in the mouse brain. One of our chief objectives was to make progress in elucidating whether the so-called ”criticality hypothesis” —in some of its possible formulations— is supported by empirical data. This goal has been tackled in two steps.

The first step was to assess the presence or absence of scale invariance or ”scaling” in neural data, for which we extensively rely on the phenomenological RG approach recently proposed by Meshulam *e*t al. [45, 46, 48]: our analyses here confirm the existence of strong signatures of scaling in all the analyzed brain regions, with exponent values taking quasi-universal values with only relatively small variations across areas [47]. The level of universality is not as precise as in critical phenomena i.e. in ”non-living matter” (though, even in Physics, critical exponents can take non-universal, continuously-varying values depending, for instance, on structural heterogeneity/disorder (see e.g. [27]). Further work is needed to better understand the origin and functional meaning of this variability.

The second step was to scrutinize whether the empirically observed scale invariance stems from criticality or not. Thus, the next sections are devoted to the discussion of this hypothesis at the light of our presented results.

### Criticality versus latent dynamical variables

Importantly, as already mentioned, recent works [50, 49] have shed doubts on the possible relation between the empirically-found scaling in neural recordings and actual criticality, as originally suggested by Meshulam *e*t al. [45, 46]. In particular, Morrell *e*t al. constructed a very simple model of binary neurons which, being uncoupled, cannot possibly exhibit collective behavior such as phase transitions or criticality [49]. In their setup, all individual neurons are exposed to a large set of shared external, time-correlated inputs, so-called ”latent dynamical variables” or simply ”hidden variables”. Surprisingly enough, such a toy model is able to reproduce —for a relatively broad region of the parameter space— the RG scaling with non-Gaussian activity distributions as well as a set of exponents roughly compatible with those in actual neural networks [45, 46]. This suggests that the empirically observed scale invariance in neural recordings could possibly have little to do with intrinsic reverberating critical dynamics, but it can emerge as an evoked response to shared external drivings.

Let us recall that this type of dichotomy for the interpretation of scaling —between critical behavior on the one hand and a superposition of the simpler effects stemming from hidden variables on the other— is a common theme in different branches of science, being for example at the roots of discussions about the meaning of the Zipf’s law in neural data [22, 58, 59, 26].

Here, by employing a variety of tools, we have concluded that all the empirically analyzed brain regions lie —to a greater or lesser extent— at the edge of instability, i.e. in the vicinity of a critical point separating stable from linearly unstable phases. Moreover, we have also shown that a random network of linear units tuned close to the edge of instability suffices to reproduce non-trivial scaling features, with remarkably similar exponent values without the explicit need of external fields.

Let us caution, however, that the previous results do not imply that external (latent) dynamical inputs may not have an impact on the recurrent dynamics nor on the observed scaling exponents. External inputs contribute to set the network working point at which covariances (as well as the stability matrix of the linear-rate model) are computed. Furthermore, the observed differences in exponent values across regions could stem from diverse exposures to latent fields from other areas.

Further empirical and theoretical studies would be required to advance in this direction; in particular, developing more elaborated models including explicitly both, near-critical recurrent dynamics and shared latent variables, as well as analyzing how the relative weights of both contributions shape scaling exponents remain open goals for future research.

Finally, a particularly relevant open question is whether the observed differences across regions can be related to the specific functional role of each area. Even if we do not have a clean-cut answer to this, let us make the following observation. One parsimonious measure of the role of a (mouse) brain area is its hierarchical score as recently determined [60], with low (high) scores corresponding to sensory (higher-level) areas. Results in Fig.4D allow us to observe that more sensory areas such as primary cortices MOp and SSp (with low scores; see e.g. Fig.6 in [60]) tend to operate closer to the edge of instability than secondary cortices (such as MOs) or areas in the prefrontal cortex (such as ORB, PL and ACA). This seems to suggest that there could exist a relationship between the dynamical regime of a given area and itsu hierarchical score, with low-score regions being more ”critical”. On the other hand, we do not observe a clear and consistent trend in thalamic regions such as LGd, LP, MG and PO. Thus, more thorough and comprehensive studies would be needed to draw solid conclusions on this inspiring possible connections.

### Edge of instability and optimal representations

By applying the phenomenological RG procedure to the data for VISp from Stringer *e*t al. [6], we found that the overall activity exhibits clear signatures of scale invariance and shares its quasi-universality with the previously analyzed regions relying on a different dataset, solving a seeming contradiction between our own results for quasi-universal scaling across brain regions and those in [6]. However, the projection of the overall or ”raw” activity into input-representing and complementary/orthogonal-space activities allowed us to conclude that the scaling exponent determined from RG analyses in the first case obeys the mathematical constraints derived by Stringer *e*t al. Understanding how the brain performs this task, i.e. how it separates signal from noise is an open fundamental problem (see e.g. [61]).

Finally, let us emphasize that, given that all the analyzed regions are near critical, they are bound to exhibit power-law decaying covariance-matrix spectra. Thus, we conjecture that critical behavior creates the broad range of covariance scales, needed for neural networks to support optimal input representations with power-law decaying eigenvalues. More research is needed to confirm this conjecture and put it on firmer ground.

To further illustrate the possible relationship between power-laws in the spectra of covariance matrices and optimal input representations, let us mention that, in a related work, we have recently designed and analyzed a simple machine-learning model based on the paradigm of *reservoir computing* [62] to analyze this problem in a well-controlled example. This model consists of a recurrent network of coupled units/neurons, which receive shared external inputs, giving rise to reverberating activity within the network or ”reservoir”. Contrarily to other machine-learning paradigms, the internal synaptic weights remain fixed during the training process: only a smaller subset of links connecting to a set of readout nodes change during training, which makes reservoir-computing a versatile tool for diverse computational tasks [63]. Inspired by the experiments of Stringer *e*t al., we trained the network on an image classification task. We refer the interested reader to [62] for further details. For our purposes here, it suffices to recall that the best performance is obtained when the tunable control parameters are set in such a way that the overall dynamical state is very close to, but below, the edge of instability. Moreover, within such an operational regime, the spectrum of the covariance matrix obeys the mathematical requirement for optimal representations, i.e., *µ ≳* 1, as observed for actual neural networks [6].

We find quite suggestive that such a relatively-simple artificial neural network becomes optimal in a regime that shares crucial statistical properties of the covariances with actual neural networks in the (mouse) brain. Thus, we believe that this machine-learning model may constitute a well-controlled starting point to further investigate the interplay between internal dynamics and external shared inputs in more realistic models of brain activity and to scrutinize optimal input representations. In any case, this seems to confirm that scale-invariant covariance spectra, as a naturally emerging property of critical systems, constitute an excellent breeding ground for optimal information storage.

### Avalanche criticality versus edge-of-instability

As already discussed, we have found strong empirical evidence of critical behavior in the sense of vicinity to the edge of instability across brain regions. This type of behavior —called traditionally ”edge of chaos” or ”type-II criticality” in [34]— has long been (since the pioneering works of Langton and others [64, 65]) theoretically conjectured to be crucial for information processing in natural and artificial neural networks. In this case, edge-of-instability systems are characterized by the presence of many modes that are close to become unstable, and thus there is a large repertoire of possible dynamical states that can be excited, opening many possible channels to information processing and transmission in real time [65, 66, 11].

However, in the analyzed neural recordings there are not large fluctuations in the overall level of global activity across time, in agreement with what observed for the motor cortex of awake macaque monkeys in [34] and with the expectation for ”asynchronous states” in balanced networks [55]. Nevertheless, it is noteworthy, that in some other empirical observations diverse levels of temporal variability in collective firing rates (e.g. gamma oscillations) have been also reported together with the possibility of bursts and avalanching behavior [7, 35, 37]. Actually, as stated in the Introduction, much attention has been paid to ”avalanche criticality” (referred as ”type-I criticality” in [34]), which is associated with networks in which there is an overall-activity mode about to become unstable, thus generating scale-free avalanches of activity, while the rest of modes are stable. Importantly, scale-free avalanches can also occur at the edge of synchronization phase transitions where the dominant mode becomes oscillatory [21, 29, 67].

It should be emphasized that both types of criticality are not mutually exclusive as, in principle, it is possible to have a whole set of eigenvalues, including the overall-activity one, at the edge of instability. Similarly, the network could be at the edge of becoming collectively oscillatory (as recently proposed for large-scale brain networks in e.g. [21, 29]) if the leading eigenvalue crosses the instability threshold with a non-vanishing imaginary part and at the same time have many other modes near the edge of instability as found here. As an illustrative example, let us mention that a computational model of spiking neurons, including both ”asynchronous states” and a synchronization phase transition characterized by scale-free avalanches, has been recently proposed and analyzed [29]. Thus, it seems a priori possible to construct computational models exhibiting both types of criticality in which the system can shift between different regimes depending on its needs. For instance, asynchronous states near the edge-of-instability support local computational tasks, while emerging synchronous oscillations are useful for information transfer and reliable communication with distant areas [68]. In our view, it is likely that actual brain networks exploit these two complementary ways of being critical to achieve diverse functional advantages for different tasks, possibly by shifting their dynamical regimes in response to stimuli.

## Conclusions

We have developed a synergistic framework which relies on recently proposed breakingthrough approaches, but that also extends and combines them to analyze state-of-the-art recordings of the activity of many neurons across brain regions in the mouse. We find that all regions exhibit scale invariance and that all of them operate, to a greater or lesser extent, in a critical regime at the edge of instability. Moreover, we have shown that the resulting scaling in the spectrum of covariances can have very important functional applications for information storage, as it facilitates the generation of optimal input representations. It is our hope that the present work stimulates further research on the remaining open questions and helps advance towards a more comprehensive understanding of the overall dynamics of brain networks and their emerging computational properties, as well as to disentangle universal and non-universal aspects across regions and behavioral states.

## Appendix

### Phenomenological renormalization-group (RG) approach

Let us briefly outline the phenomenological RG approach introduced in [45] and [46]. Given a set of *N* neurons, the empirically-determined activity of the *i*-th neuron at a given time *t*_*j*_ is denoted by *σ*_*i*_(*t*_*j*_), where *j* ∈ (1, *T*) labels a discrete number of non-overlapping time bins. Determining the most meaningful size of such time bins is an important technical aspect (see next section); here, we just assume that such an optimal time discretization is given.

As discussed in the main text, a criterion of maximal pairwise correlation is employed to group neurons together at each step *k* of the RG procedure. In particular, one considers the Pearson’s correlation coefficients

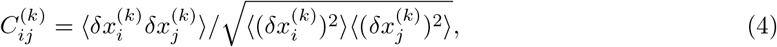

Where 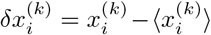 and 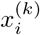 is the activity of the block-neuron *i* at step *k* of the coarse-graining (we identify 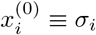 as the activity of neuron *i* before coarse-graining), while averages are computed across the available discrete time steps. At the beginning of each RG step, the two most correlated neurons, *i*, and *j*_*∗i*_, are selected and their activities are added into a new coarse-grained variable:

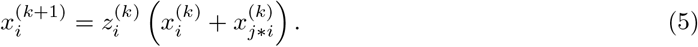

where the normalization factor 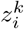 is chosen in such a way that the average non-zero activity of the new variables 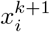 is equal to one. Notice that, owing to such a normalization criterion, activity values are not constrained to fulfill 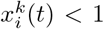. Then one proceeds with the second most-correlated pair of neurons and so on, until a set of *N*_*k*_ coarse-grained ”block neurons”, each containing the summed activity of *K* = 2^*k*^ original neurons, has been constructed.

Iterating this procedure, after *k* steps there remain only *N*_*k*_ = *N/*2^*k*^ coarse-grained variables or ”*block-neurons*”, 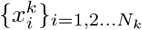, each recapitulating the activity of *K* = 2^*k*^ individual neurons. To figure out whether a fixed point of the RG flow exists, one can study the evolution of the probability density function for the activity of the coarse-grained variables, *P*_*K*_(*x*). Following [45], we separated *P*_*K*_(*x*) for block-neurons of size *K* in two components: the probability of being silent, *S*_*K*_, and the probability *Q*_*K*_(*x*) of having non-zero activity *x*: *P*_*K*_ (*x*) = *S*_*K*_*δ* (*x*) + (1 − *S*_*K*_)*Q*_*K*_ (*x*). Trivially, if the original neurons were statistically independent, one would expect (as a direct consequence of the central limit theorem) to drive the activity distribution *Q*_*K*_ towards a Gaussian fixed-point of the RG flow. As pointed out in [45], the RG convergence to a non-Gaussian fixed-point (i.e. invariance of the distribution across RG steps) reveals a non-trivial structure in the data.

Another quantity of interest is the variance of the activity distributions as a function of the size of the block neurons *K*:

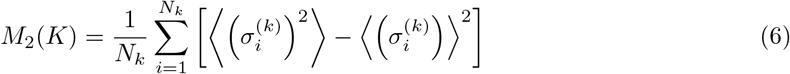

where 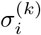 is the summed activity of the original variables inside the cluster. Notice that, for totally independent variables, one would expect the variance to grow linearly in *K* (i.e., *M*_2_(*K*) ∝ *K*), whereas if variables were perfectly correlated *M*_2_(*K*) ∝ *K*^2^. Non-trivial scaling is therefore characterized by *M*_2_(*K*) ∝ *K*^*α*^ with a certain intermediate value of the exponent 1 *< α <* 2.

On the other hand, *F*_*k*_ = − log (*S*_*K*_) defines a sort of ”free-energy” for the coarse-grained variables at the *k*-th RG step [45]. As more and more of the initial variables *σ*_*i*_ are grouped into cluster variables 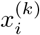, one would expect that the probability of having ”silent” block-neurons (i.e., the probability that all neurons inside a cluster are silent) decreases exponentially with the size *K* of the clusters, leading to: *F* (*K*) ∝ *K*^*β*^ where *β* = 1 for initially independent variables.

One can also wonder whether there is some type of self-similarity in the dynamics at coarse-grained scales. Given that, commonly, fluctuations on larger spatial scales relax with a slower characteristic time scale, we should expect the time-lagged Pearson’s correlation function (or simply autocorrelation function) of the coarse-grained variables to decay more slowly as we average over more neurons. In particular, for step *k* of the RG flow, one has:

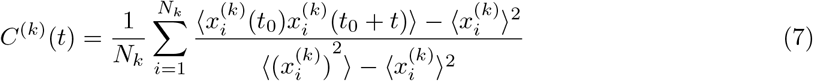

Assuming that correlations decay exponentially in time with a characteristic time scale 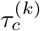 (i.e., 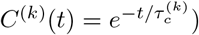) at each coarse-graining level, dynamical scaling implies that the average correlation function collapses into a single curve when time is re-scaled by the characteristic time scale: 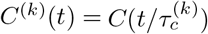 and that this time scale obeys scaling with the cluster size: *τ* (*K*) ∝ *K*^*z*^ where *z* is the dynamical scaling exponent. Finally, as argued in [45], if correlations are self-similar then we should see this by looking inside the clusters of size *K*. In particular, the eigenvalues of the covariance matrix (i.e., the propagator, which is scale-invariant at the fixed-point of the RG in systems with translational invariance [45]) must obey a power-law dependence on the fractional rank: *λ* = *B* (*K/rank*)^*µ*^, where *B* is a constant and *µ* a decay exponent.

We estimate the goodness of each power-law fit by calculating the R-squared value, comparing too with an equivalent exponential fit (see Fig.S9 in the SI) [69]. For the probability density of eigenvalues we compute the log-likelihood ratios between the estimated power-law and alternative exponential and lognormal distributions (see Extended Methods and Tables S1 and S2 in the SI).

Each exponent is expressed as 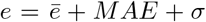 + *MAE* + *σ*, where 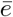 is the average across different experiments (possibly from different mice), *MAE* is the mean-absolute-error, computed as the average across experiments of the experiment-specific errors measured over split-quarters of data, and *σ* is the standard deviation across experiments (see Tables S1 and S2 in SI).

### Measuring covariances

Here, we use a general definition of covariance, as described by the following equation:

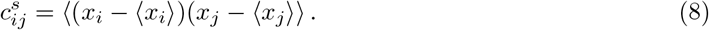

During the RG analysis and in estimations of the distance to the edge of instability, the variable *x*_*i*_ represents the number of spikes in a time bin of width Δ*t*, with averages taken over all timebins, so that *c*_*ij*_ measures the pairwise correlation of spike-count responses to repeated presentations of the same stimulus (or, in our case, repeated sampling of resting-state activity under identical behavioral conditions). Throughout this article, we referred simply as ”covariance” when correlations were computed in the RG analysis using the time bin given by the geometric mean of neurons’ ISIs, which renders time series where the average non-zero bin is populated by only one spike. In contrast, the term ”long-time-window” or ”spike-count” covariance is left for the distance to criticality analysis, in which Δ*t* = 1s and the average non-zero bin in the time-series contains between 3 and 7 spikes, depending on the region. We notice that the later can be written as the time integral of the time-lagged covariance *c*_*ij*_(*τ*) [34]:

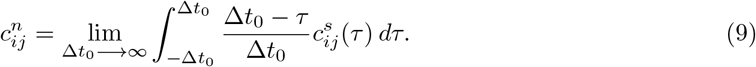

which, loosely speaking, removes time-dependent effects and puts the emphasis onto pairwise heterogeneities.

### Extended Methods

See the ”Extended Methods” section in the Supplementary Information text for additional details.

## Acknowledgements

We acknowledge the Spanish Ministry and Agencia Estatal de investigación (AEI) through Project of I+D+i Ref. PID2020-113681GB-I00, financed by MICIN/AEI/10.13039/501100011033 and FEDER “A way to make Europe”, as well as the Consejería de Conocimiento, Investigación Universidad, Junta de Andalucía and European Regional Development Fund, Project references A-FQM-175-UGR18 and P20-00173 for financial support. We also thank V. Buendía, P.Villegas, R. Corral, J. Pretel, P. Moretti, M. Ibañez, P. Garrido, and M. Marsili, for valuable discussions and/or suggestions on earlier versions of the manuscript.

