## Supplementary Information for "Quasi-universal scaling in mouse-brain neuronal activity stems from edge-of-instability critical dynamics"

[a] *Departamento de Electromagnetismo y Física de la Materia and Instituto Carlos I de Física Teórica y Computacional. Universidad de Granada. E-18071, Granada, Spain*
[b] *Center for Theoretical Neuroscience, M. B. Zuckerman Mind Brain Behavior Institute, Columbia University, New York, NY USA*

### 1. Extended Methods

#### Datasets

In the experimental setup of Steinmetz *e*t al., mice were exposed to visual stimuli of varying contrast that could appear on the left side, right side, both sides or neither side of a screen. Mice then earned a water reward by turning or holding still a wheel, depending on the type of stimuli. During task performance, two or three Neuropixels probes were inserted at the same time in the left hemisphere, allowing for high-resolution simultaneous-recordings of hundreds of neurons in several regions during each recording session [1]. Thus, for each session the dataset provides the time-stamps and amplitudes of all measured spikes within various brain regions for a total recorded time $T$. In our analysis, we first collected all spikes belonging to a given region, provided it contained more than 128 simultaneously recorded neurons. Then, an optimal time bin $\Delta t$ was inferred for each area on the basis of the Inter-Spike-Interval distribution of its neurons (see *Time-scale determination* below). This allowed us to convert the spike time-stamps of each neuron into discrete trains of spikes, which were later used to infer pair-wise correlations among neurons and, ultimately, perform RG analyses.

On the other hand, in the work of Stringer *e*t al. mice were presented a sequence of 2800 images, each remaining for $0.5s$ and intersperse with gray-screen inter-stimuli intervals lasting a random time between $0.3s$ and $1.1s$ [2]. The resulting task-related activity is thus effectively binned into $\Delta t \approx 0.5s$ time intervals (a time resolution that we keep in our analyses) and two repetitions of the same input-presentation sequence were performed to allow for cv-PCA analyses. Experiments were then repeated with 3 different mice and using different dimensions of the input ensemble. To construct d-dimensional stimuli, the natural image database was projected onto a set of basis functions that enforced the required dimensionality [2]. In our analyses we show results for natural, high-dimensional images as well as 4-d, low-dimensional images.

#### Time-scale determination

In order to determine pairwise correlations from empirical neural activity data, it is necessary to discretize the time into bins of certain length $\Delta t$. In our case, to obtain a spike train vector from a list of time stamps $t_j^*$ at which a neuron spikes, a common value $\Delta t$ is chosen for all units. In this way, neurons spiking at times $t_j$ and $t_j + \delta t$ sufficiently close share a bin $\Delta t_j$ with non-zero activity. The number of bins for a neuron can be therefore chosen by $T_i = \frac{max(t_j^*)}{\Delta t}$, and more generally we will

set $T = max(T_i)$ so that all neurons have the same amount of bins in their spike trains.

Choosing the time bin that best transforms the spike times into discrete trains of spikes is an open problem in neuroscience for which several solutions —such as the use of random bins [3], methods that find the best bin size by minimizing a certain cost function [4, 5, 6, 7] and bin-less approaches [8, 9] have been proposed. The problem becomes even more prominent when working with the simultaneously-recorded activity of a large number of cells as, typically, one finds neurons operating at broadly different time scales within the same area (see Fig.3), as well as a hierarchy of timescales across regions [10, 11].

To perform the RG analyses, we first compute the distribution of inter-spike intervals (ISI) of individual neurons within a particular brain region (Fig.4). Because we are interested in a measure of the "typical" time-scale at which neurons in a population operate, and given also the large neuron-to-neuron variability, we define the optimal time bin $\Delta t$ as the geometric mean of all ISIs from neurons in the population which (as illustrated in Fig.4), is a very good estimator of the population characteristic time scale); this geometric-mean value is computed for each region and subsequently used in the corresponding RG analysis (see Table 1 for a summary of all areas with their corresponding selected time bin).

#### Stationarity of spiking statistics

The expression for the maximum eigenvalue $\lambda_{max}$ of the effective connectivity matrix (as derived originally by Dahmen et al. using linear response theory) relies on the stationarity of the spike trains. Therefore, to assess the stationarity of the recordings in each region we performed an Augmented Dickey-Fuller test (ADF) [12], which evaluates the null hypothesis that a unit root is present in a time series (i.e., it has some time-dependent structure and it is therefore non-stationary). The alternative hypothesis, should the null hypothesis be rejected, is that the time series is stationary. The *adfuller* function from the *statsmodel* Python library was used to perform the analysis.

Thus, for each neuron in a particular region and experiment we tested the null hypothesis at a 5% significance level, then counted the fraction of neurons in the experiment for which the null hypothesis could not be rejected (i.e. had non-stationary activity). This fraction was then averaged over all experiments to estimate the percentage of neurons with non-stationary activity for a given region (see Table 1), and only those experiments with less than 10% of the neurons showing non-stationary spiking statistics were considered in the subsequent analyses.

#### Autocorrelation decay times

Spike-trains were first smoothed using a Gaussian-kernel with $\sigma = 50ms$ to obtain a firing rate time-series. For each neuron, the autocorrelation function was then computed, averaging across all neurons to obtain an estimate of the autocorrelation for the population (reported for the MOp region as an example in Fig.4A of the main text). We then fitted the resulting decaying function to an exponential, extracting a characteristic time-scale $\tau_{Corr}$ for each experiment. Finally, these values were averaged across experiments to obtain the mean autocorrelation decay time in each region (see Table 1). The results allow us to define an optimal $\Delta t = 1s$ over which to compute spike-counts covariances, sufficient for autocorrelations to decay (a requirement for the equivalence between integrated time-lagged and spike-count covariances to hold) while maximizing the amount of available samples.

#### Likelihood ratio test for powerlaw distribution of eigenvalues

In the limit of $N \to \infty$ there exist a direct relationship between the exponent of the rank-ordered eigenvalues and their probability density (see Section 5 below). Therefore, we leveraged the approach in [13] to test whether a power-law distribution is indeed the best description of the density of eigenvalues for the different regions in the data set. Following [13] we computed the loglikelihood ratio $R$ between the estimated power-law and equivalent exponential and lognormal candidate distributions, together with the the p-value for the significance of the test. In 14 out of the 16 regions, the exponential distribution provides a worse fit —tested at a 5% significance level— than the corresponding power-law ($R > 0$ ; $p \leq 0.05$). On the other hand, the lognormal distribution always provides an equally good description of the eigenvalue density ($p > 0.05$), and cannot be ruled out as the underlying distribution. These values, together with the R-squared value for the best powerlaw and exponential fits of the rank-ordered eigenvalues in each region, are listed in Table 2, whereas the corresponding fitted curves are plotted in Fig.9.

##### Estimation of parameter g

The dynamics of a neuron in a recurrent network model of linear units driven by noise is described by the following equation:

$$\dot{x_i}(t) = -x_i(t) + g\sum_{i=1}^{N} J_{ij}x_j + \xi_i(t), \quad i = 1, ..., N \tag{1}$$

with i.i.d. Gaussian connectivity $J_{ij} \sim \mathcal{N}(0, 1/N)$ and $\langle \xi_i(t)\xi_j(t+\phi)\rangle = \delta_{ij}\delta(\phi)$. For this type of model, the dynamics becomes unstable at a critical value $g_c = 1$ [14], where $g$ describes the overall connection strength (i.e., the spectral radius or maximum eigenvalue of the connectivity matrix, $\lambda_{max}$, as defined in [15]) and can be considered a control parameter of the system. In the large $N$ limit the analytical form of the probability density of the covariance eigenvalues has been derived in [14], and reads:

$$p_{RC}(x) = \frac{3^{\frac{1}{6}}}{2\pi g^2 x^2}\left[\sum_{\xi=1,-1}\xi\left((1+\frac{g^2}{2})x - \frac{1}{9} + \xi\sqrt{\frac{(1-g^2)^3 x(x_+ - x)(x - x_-)}{3}}\right)^{\frac{1}{3}}\right], \quad x_- \leq x \leq x_+, \tag{2}$$

with

$$x_{\pm} = \frac{2 + 5g^2 - \frac{g^4}{4} \pm \frac{1}{4}g(8+g^2)^{\frac{3}{2}}}{\cdot} \tag{3}$$

Thus, given a data set of simultaneously recorded neurons, one can infer the dynamical regime of the population activity by fitting the sampled covariance matrix eigenvalue spectrum to the theoretical distribution. More concretely, we looked for the value of $g$ that minimized the $L^2$ error using the Cramer-von Mises statistics between the empirical cumulative distribution $F_n(x)$ and the theoretical one $F(x) = \int_{-\infty}^{x} p_{RC}(x)dx$ [14]:

$$D^2_{CvM} = \int (F(x) - F_n(x))^2 dF_n(x) = \frac{1}{12n^2} + \frac{1}{n}\sum_{i=1}^{n}\left(F(x_i) - \frac{2i-1}{2n}\right), \tag{4}$$

where $n$ is the total number of samples and $x_i$ are the eigenvalues of the empirical long time window covariance matrix. For a given region, we computed $D^2_{CvM}$ for each experiment, then averaged across all experiments (values are provided in Table 1).

#### 2. Further results of the RG analyses

Here we provide extended results illustrating the ubiquity of scaling across brain regions.

##### Non-zero activity distributions across brain regions

Fig.1 shows the normalized probability distribution function for the non-zero activity inside clusters of size $K$ (and averaged over all existent clusters) at 5 different steps of the coarse-graining. As in the main text, errors are computed as the standard deviation over random quarters of the data, and only shown in the last step of coarse-graining.

#### Scaling of time-decaying auto-correlation curves

Fig.2 shows the auto-correlation function for resting-state type of activity, computed as:

$$C^{(k)}(t) = \frac{1}{N_k} \sum_{i=1}^{N_k} \frac{\langle x_i^{(k)}(t_0) x_i^{(k)}(t_0+t)\rangle - \langle x_i^{(k)}\rangle^2}{\langle \left(x_i^{(k)}\right)^2 \rangle - \langle x_i^{(k)}\rangle^2} \tag{5}$$

and normalized to lie within the unit interval. In each subplot, the above quantity is plotted against the re-scaled time $t/\tau_c$, where $\tau_c$ is the characteristic time of the exponential decay, for a particular region of the mouse brain at for 4 different levels of coarse-graining. For ease of visualization, the errors —computed as the standard deviation over random (non-shuffled) quarters of the data— are only shown in the last step of coarse-graining procedure.

#### Robustness against time bin choice

Activity of neurons, even within the same brain region, can span many time scales, with some neurons typically firing in the range of milliseconds and with a characteristic time-scale of the order of seconds (see Fig.3). Thus, in selecting a common time bin to convert the spike times of all neurons within a region into discrete trains of spikes, we irretrievably lose some information.

To assess whether the exponent values reported in the main text are robust against our choice of $\Delta t$, we repeated the same RG analyses using different time-windows $\Delta t \in [0.01s, 4s]$ to bin the resting-state activity of each region.

As illustrated in Fig.5, the values of the exponents $\alpha$, $\beta$ and $z$ turn out to be fairly robust within a broad range of biologically plausible time scales. As one could trivially expect, broader time bins increase the probability of finding neurons spiking simultaneously, i.e. appearing as more strongly correlated. This, in turn, has the effect of shifting the values of the exponents $\alpha$ and $\beta$ for the variance and free-energy scalings slightly towards those expected for a fully correlated scenario. On the other hand, the exponent $\mu$ remains fairly unchanged for binning times up to $\sim 100ms$, but then increases on longer time scales beyond the typical geometric mean ISI of the regions.

#### Resting-state vs task-induced activity

We plotted in Fig.6 the four measured scaling exponents in task-induced vs resting-state type of activity, with results that clearly support the idea that the observed exponents are almost invariant with the mouse behavioral state. Only small deviations are observed in a few regions.

#### Non-universal $\alpha$ exponent in input-encoding subspace

Fig.7 shows the scaling exponent for the variance of the activity inside block-neurons, measured over the overall raw data, the input-related/encoding activity, and the background activity. We only observe a significant change on the exponent —for both types of stimuli: natural and 4D images— when the input-encoding activity is considered, but not in the background, orthogonal subspace. Observe also, that there is no significant change in the $\alpha$ exponent in orthogonal components as compared to raw/overall activity.

#### Dependence of $\mu$ with the samples-to-neurons ratio

The spectrum of covariance eigenvalues is known to be strongly sensitive to the ratio $T/N$, where $N$ is the total number of neurons recorded in the experiment and $T$ is the number of time bins [16].

In Fig.8 we scrutinize the dependence of the scaling exponent for the rank-ordered eigenvalues of the covariance matrix $\mu$ with the ratio $a_0 = T/N$. The figure shows that the estimates converge to the asymptotic value only for sufficiently large values $a_0 > 20$, while $\mu$ typically increases as one moves towards the sub-sampled regime.

In the inset of Fig.8 we compare the fitted exponents for the rank-ordered plot of eigenvalues in the example region MOp when i) taking the activity recorded from each of the 4 intervals of spontaneous activity in the recording session, ii) merging the 4 intervals together and iii) averaging over quarter splits of the data set where the intervals were merged together. In individual spontaneous intervals $a_0 > 20$ is not always fulfilled, thus causing variability in the fitted exponents. By merging together all recordings of spontaneous activity belonging to the same experiment we can typically ensure that $a_0 > 20$, even if analyses are performed over quarter-splits of data. This confirms the consistency and robustness of the estimated exponents.

#### 3. RG analyses on surrogated data

As a sanity check, we performed the RG analysis over surrogated data in three different ways: i) by randomly shuffling all the spikes for each neuron; ii) by shifting all the spikes of each neuron a random amount, so that the structure within the spike train is preserved; and iii) by randomizing the spikes across neurons but not time. Here, we chose the MOp region for having the greatest number of recorded neurons, and compare the results with the same analysis in the unsurrogated case (Fig.11)

We observe that time series shuffle (Fig.12) or shifting (Fig.13) destroys the non-trivial scaling properties, showing no signs of convergence towards a non-Gaussian fixed point of the probability distribution, and exponents $\alpha$, $\beta$ and $z$ akin to those expected for an independent-neurons model. The power-law spectrum of the clusters covariance matrix still seems to decay with a smaller, non-trivial exponent, probably due to the firing rate heterogeneity in the population activity, but closer inspection reveals that a power-law dependence is no longer suitable when compared to the unsurrogated case.

#### 4. RG analyses on a randomly recurrent network of linear rate neurons

We simulate a randomly connected recurrent network of linear rate neurons driven by noise described by Eq.1. The simulations were ran for networks of size $N = 1024$, choosing an integration step $h = 0.01$ while setting $\sigma = 1$. We then binned the resulting time series with a $\Delta t = 0.1$ sampling window, and performed the RG analyses for various values of the control parameter $g$ (Fig.10). In particular, we show the scaling of the variance, autocorrelation time, and covariance eigenvalues inside the clusters formed along the RG flow. We observe that power-law scaling (with exponent values $\alpha$ and $z$ similar to the empirically measured ones) can be retrieved only when the system is close to the edge-of-instability ($g = 0.95$). Away from such a regime one finds the trivial expectations for asymptotically uncorrelated variables ($\alpha = 1$, $z \approx 0$). As for the covariance-matrix spectrum, we see an increasing overlap of the rank-ordered spectrum across RG steps and a tendency towards scaling collapse and power-law-like behavior as the parameters gets closer to criticality. Since the activity of the neurons in the model take positive and negative values within a continuous range, the exponent $\beta$ for the scaling of the probability of being silent is trivially zero (i.e. this property cannot be described by the simple linear model).

#### 5. Rank-ordering and spectral density of covariance matrix eigenvalues

Let us provide a simple mathematical calculation that highlights the relationship between the power-law exponents for the probability distribution of eigenvalues, that we also call "spectral density", and eigenvalue-vs-rank plots.

Let us assume that one has measured the spectrum of a covariance matrix extracted from some experimental data and found that the $n$-th largest eigenvalue scales with its rank (when ordered from the largest to the smallest) as:

$$\lambda_n = An^{-\gamma}. \tag{6}$$

Given such a dependency, what can be said about the spectral density $\rho(\lambda)$, which describes the probability of finding a certain value of $\lambda$ in the spectrum? Formally, and since there is a discrete number of eigenvalues (limited by the number of variables $N$):

$$\rho(\lambda) = \frac{1}{N}\sum_n \delta\left(\lambda - \lambda_n\right). \tag{7}$$

Nevertheless, in the limit of $N \to \infty$ one can approximate the above distribution by a continuous one:

$$\rho(\lambda) \approx \frac{1}{N}\int_1^\infty \delta\left(\lambda - \lambda_x\right) dx. \tag{8}$$

Let us now define the function $g(x) = \lambda - \lambda_x = \lambda - Ax^{-\gamma}$, which has a root at $x_0 = (\lambda/A)^{-1/\gamma}$. Using the properties of the Dirac delta distribution:

$$\delta\left(g(x)\right) = \frac{\delta\left(x - x_0\right)}{\left|g'\left(x_0\right)\right|} = \frac{\delta\left(x - x_0\right)}{\nu A\left(\frac{\lambda}{A}\right)^{1+1/\gamma}} \tag{9}$$

one can then, finally, write the spectral density in continuous approximation as:

$$\rho(\lambda) \approx \frac{1}{N}\left(\nu A\left(\frac{\lambda}{A}\right)^{1+1/\gamma}\right)^{-1}\int_1^\infty \delta\left(x - x_0\right) dx = \frac{A^{1/\nu}}{N\nu}\lambda^{-1-1/\gamma} \tag{10}$$

Thus, we can conclude that given a power-law dependency $\lambda_n \sim n^{-\gamma}$ of the eigenvalues on their rank, its corresponding spectral density is expected to also follow a power-law $\rho(\lambda) \sim \lambda^{-\nu}$, with $\nu = 1+1/\gamma$. For a more detailed discussion of this scaling relation see [17, 18].

### 6. cvPCA: noise vs stimuli-related activity

Following Stringer *et al.*, let us consider two observation matrices, $X_{(1)}, X_{(2)} \in \mathbb{R}^{N\times T}$, corresponding to two identical realizations or trials of the experiment, with $N$ the number of neurons and $T$ the number of images or stimuli presented. Thus, we denote by $x_{ij}^{(k)}$ the matrix element containing the activity of neuron $i$ during presentation of image $j$ in trial $k$. For simplicity, we will further assume that the mean activity of each neuron across images has been subtracted, so that both matrices have zero-mean rows. For each matrix one can then define two possible covariance matrices:

$$C_u = \frac{1}{T-1}XX^T \Longrightarrow C_u\mathbf{u}_i = \lambda_i\mathbf{u}_i \tag{11}$$

$$C_v = \frac{1}{N-1}X^TX \Longrightarrow C_v\mathbf{v}_i = \lambda_i\mathbf{v}_i \tag{12}$$

where $\mathbf{u}_i \in \mathbb{R}^{N\times 1}$ and $\mathbf{v}_i \in \mathbb{R}^{T\times 1}$ are the eigenvectors (singular vectors) associated with each of the above defined covariance matrices, which also share the same eigenvalues $\lambda_i$. In general, there are $r = \text{rank}\,(X)$ non-zero eigenvectors of each type. The Singular Value Decomposition (SVD) theorem states that any observation matrix can be decomposed as:

$$X = USV^T \tag{13}$$

$$XV = US \Longrightarrow X\mathbf{v}_i = \sigma_i\mathbf{u}_i \tag{14}$$

where $U \in \mathbb{R}^{N \times r}$ contains the $\mathbf{u}_i$ eigenvectors by columns, $V \in \mathbb{R}^{T \times r}$ contains the $\mathbf{v}_i$ eigenvectors by columns, and $S \in \mathbb{R}^{r \times r}$ is a diagonal matrix containing the singular values $s_{ii} = \sigma_i = \sqrt{\lambda_i}$.

Assume one has performed SVD on the first trial observation matrix $X_{(1)}$, obtaining the singular vectors $U_{(1)}$ and $V_{(1)}$. Defining the projection matrix $P = U_{(1)}^T \in \mathbb{R}^{r \times N}$, which diagonalizes $C_u^{(1)} = \frac{1}{T-1} X_{(1)} X_{(1)}^T$, one can compute:interval = [0,1,3]

$$Y_{(1)} = P X_{(1)} \Longrightarrow \mathbf{y}_i^{(1)} = (\mathbf{u}_i^{(1)})^T X_{(1)} \in \mathbb{R}^{1 \times T} \tag{15}$$

where $Y_{(1)} \in \mathbb{R}^{r \times T}$ contains the projection of the first trial activity over its $r$ principal components. Now, the idea of the cv-PCA is to project on the same subspace the observations coming from the second trial [2]:

$$Y_{(2)} = P X_{(2)} \Longrightarrow \mathbf{y}_i^{(2)} = (\mathbf{u}_i^{(1)})^T X_{(2)} \in \mathbb{R}^{1 \times T} \tag{16}$$

and ask to form a covariance matrix with the product of both projections:

$$C_\psi = \frac{1}{T-1} Y_{(1)} Y_{(2)}^T = \frac{1}{T-1} \begin{pmatrix} \mathbf{y}_1^{(1)} \cdot \mathbf{y}_1^{(2)} & \mathbf{y}_1^{(1)} \cdot \mathbf{y}_2^{(2)} & \cdots \\ \mathbf{y}_2^{(1)} \cdot \mathbf{y}_1^{(2)} & \mathbf{y}_2^{(1)} \cdot \mathbf{y}_2^{(2)} & \cdots \\ \cdots & \cdots & \mathbf{y}_r^{(1)} \cdot \mathbf{y}_r^{(2)} \end{pmatrix}. \tag{17}$$

One can now hypothesize that the activity at time $t$ of any neuron $i$ during trial $k$ can be linearly decomposed as:

$$\mathbf{x}_i^{(k)}(t) = \psi_i(t) + \epsilon_i^{(k)}(t) \tag{18}$$

where $\psi_i$ denotes the input-related activity, which should be independent of the trial for experiments carried in identical conditions; and $\epsilon_i^{(k)}$ is defined as background or trial-to-trial variable activity, spanning a subspace that is orthogonal to the input-related one. One can then show that the i-th diagonal element of $C_\psi$ is a non-biased estimator of the input-related variance $\omega_i$ [2]:

$$\omega_i = \frac{1}{T-1} \mathbf{y}_i^{(1)} \cdot \mathbf{y}_i^{(2)} = \frac{1}{T-1} \left( (\mathbf{u}_i^{(1)})^T X_{(1)} \right) \left( (\mathbf{u}_i^{(1)})^T X_{(2)} \right)^T = \frac{1}{T-1} (\mathbf{u}_i^{(1)})^T X_{(1)} X_{(2)}^T \mathbf{u}_i^{(1)}. \tag{19}$$

Rewriting the observation matrices as $X_{(k)} = \Psi + \Sigma_{(k)}$, where $\Psi$ contains vectors $\psi_i$ in rows and so does $\Sigma_{(k)}$ with vectors $\epsilon_i^{(k)}$:

$$\omega_i = \frac{1}{T-1} \mathbf{u}_i^T \left( \Psi + \Sigma_{(1)} \right) \left( \Psi + \Sigma_{(2)} \right)^T \mathbf{u}_i = \frac{1}{T-1} \mathbf{u}_i^T \Psi \Psi^T \mathbf{u}_i + \frac{1}{T-1} \mathbf{u}_i^T \Sigma_{(1)} \Psi^T \mathbf{u}_i \tag{20}$$

where all the terms containing $\Sigma_{(2)}$ can be dismissed due to statistical independence of the realizations. Under reasonable approximations shown in [2], one can prove that if the singular vectors $\mathbf{u}_i$ approach the singular vectors of $\Psi$, then the first term converges to the actual variance of the input-related activity along the principal direction $\mathbf{u}_i$, while the second term converges to zero. We have thus seen how the cv-PCA method proposed by Stringer *et al.* allows one to estimate the input-related covariances.

Is it possible to take a step further and find a proxy for the input-related activity observation matrix $\Psi \in \mathbb{R}^{N \times T}$, such that $\tilde{C}_\psi = \frac{1}{T-1} \Psi \Psi^T$ has the same eigenvalue spectrum as $C_\psi$? To do so, we first define a new basis of projected vectors given by:

$$\mathbf{z}_i = \sqrt{\mathbf{y}_i^{(1)} \cdot \mathbf{y}_i^{(2)}} \frac{\mathbf{y}_i^{(2)}}{\left\| \mathbf{y}_i^{(2)} \right\|} \in \mathbb{R}^{1 \times T}. \tag{21}$$

which clearly fulfills $\mathbb{E}\left[ \mathbf{z}_i \cdot \mathbf{z}_i^T \right] = \omega_i$ under the same conditions as above. If $Z \in \mathbb{R}^{r \times T}$ is the matrix composed by the $r$ vectors $\mathbf{z}_i$ in rows, then:

$$\Psi = P^T Z. \tag{22}$$

is the input-related activity of the neurons. From there, it is straightforward using Eq.18 that one can estimate the background activity just by subtracting the input-related activity from the raw data:

$$\Sigma_{(k)} = X_{(k)} - \Psi. \tag{23}$$

Thus, following the above steps, we have found a proxy (Eq.18) to decompose the overall activity of any neuron at a certain time step $t$ into an encoding (input-related) and background contributions.

For the sake of consistency, Fig.15) shows the spectrum of the covariance matrix as estimated from such a decomposition, illustrating that, to a very good approximation it coincides with the above-computed one (using the standard cv-PCA method as discussed in [2].

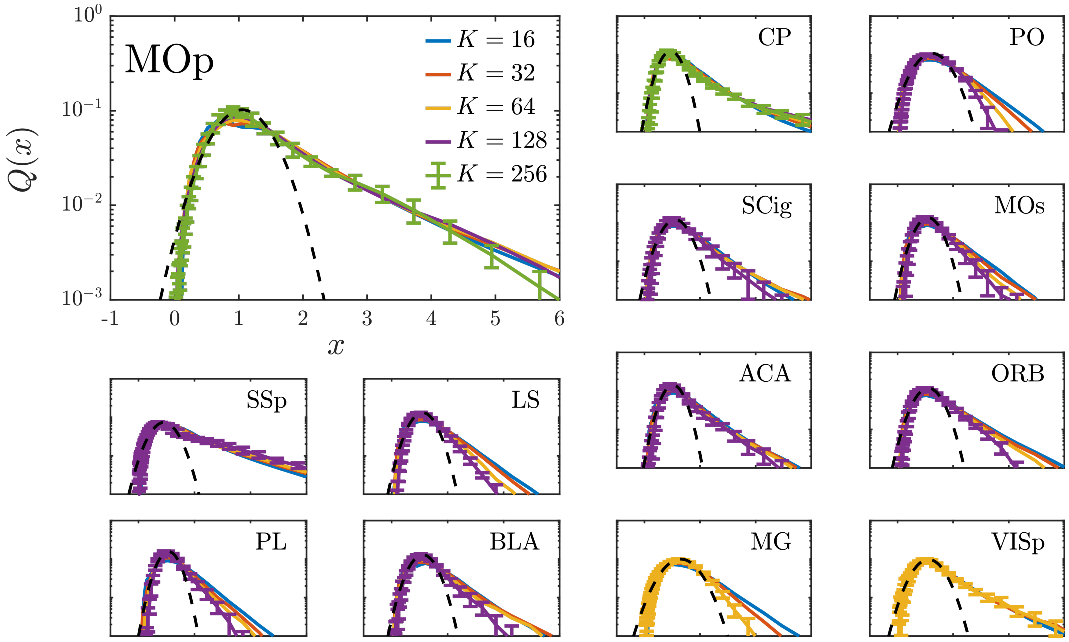


Figure 1: Non-zero activity distribution curves at different levels of coarse-graining in 13 different areas of the mouse brain as presented in the main text. Significant tails towards large values of the activity are observed, while curves are fairly invariant in most areas across RG steps, suggesting the existence of a non-Gaussian fixed point of the RG flow. Errors (shown only for the curve corresponding to the last step of coarse-graining) are computed as the standard deviation over random quarters of the data.

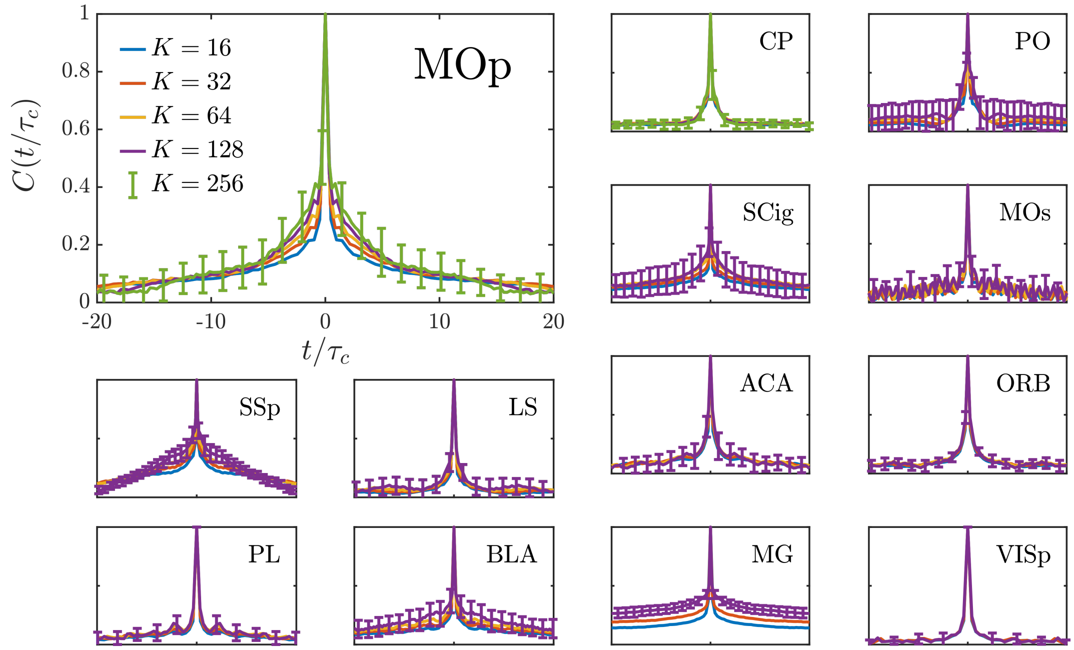


Figure 2: Mean normalized correlation function of coarse-grained variables during the RG flow, measured in 13 different areas of the mouse brain. Time is re-scaled for each curve by the characteristic time scale $\tau_c(K)$, computed as the $1/e$ point of the decay. Errors shown in the last step of coarse-graining are computed as the standard deviation over random quarters of the data

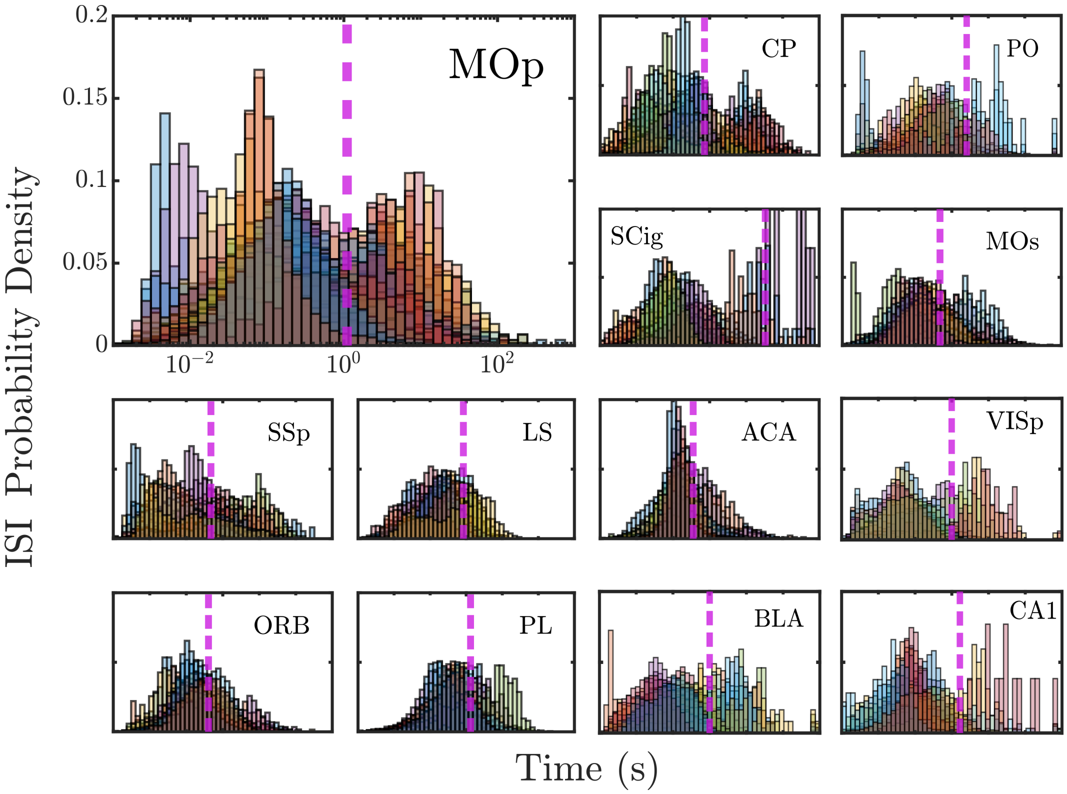


Figure 3: Histograms estimating the probability of finding a certain value of the inter-spike interval (ISI) in the resting-state activity of individual neurons belonging to a particular area (shown, as before, for 13 areas). Each color represents the distribution for a single neuron (only a few neurons are shown for the sake of clarity), while pink dashed lines mark the geometric mean ISI for each area. Let us remark that the arithmetic mean does not properly represent the characteristic time scale in each region.

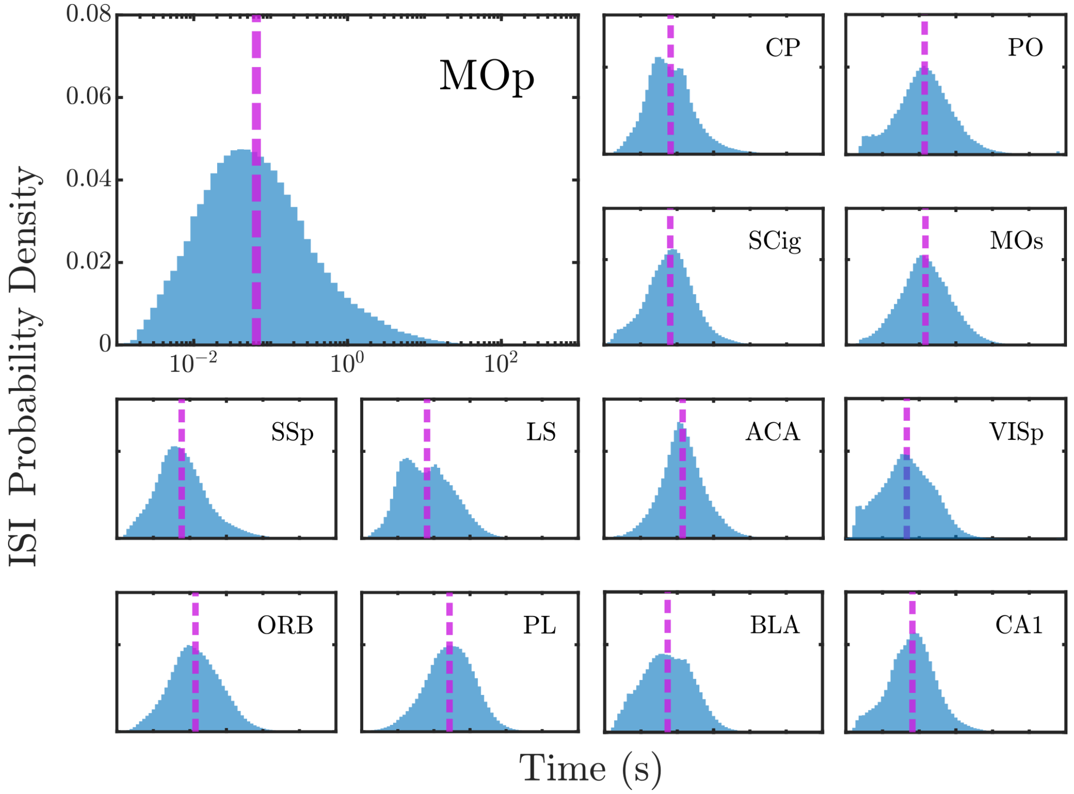


Figure 4: Individual-neuron inter-spike interval distributions for 13 different regions of the mouse brain as determined using all recorded neurons in each population (for each of them, there is a large neuron-to-neuron variability as shown in Fig.3). The characteristic or ”optimal” time bin is defined as the geometric mean of the ISI distribution (vertical dashed lines).

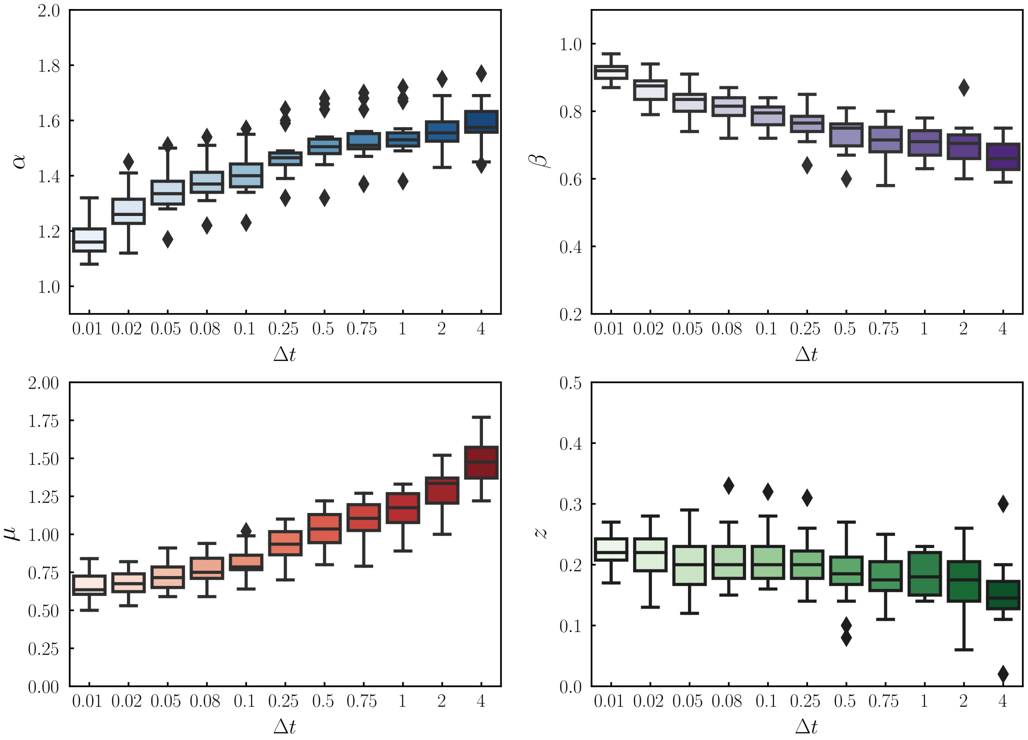


Figure 5: Change of average scaling exponents as a function of the employed time bin within the range $\Delta t \in [0.01s, 4s]$ for each region. The line inside of each box represents the sample median, the whiskers reach the non-outlier maximum and minimum values; the distance between the top (upper quartile) and bottom (lower quartile) edges of each box is the inter-quartile range (IQR). Black diamonds represent outliers, i.e. values that are more than $1.5IQR$ away from the limits of the box.

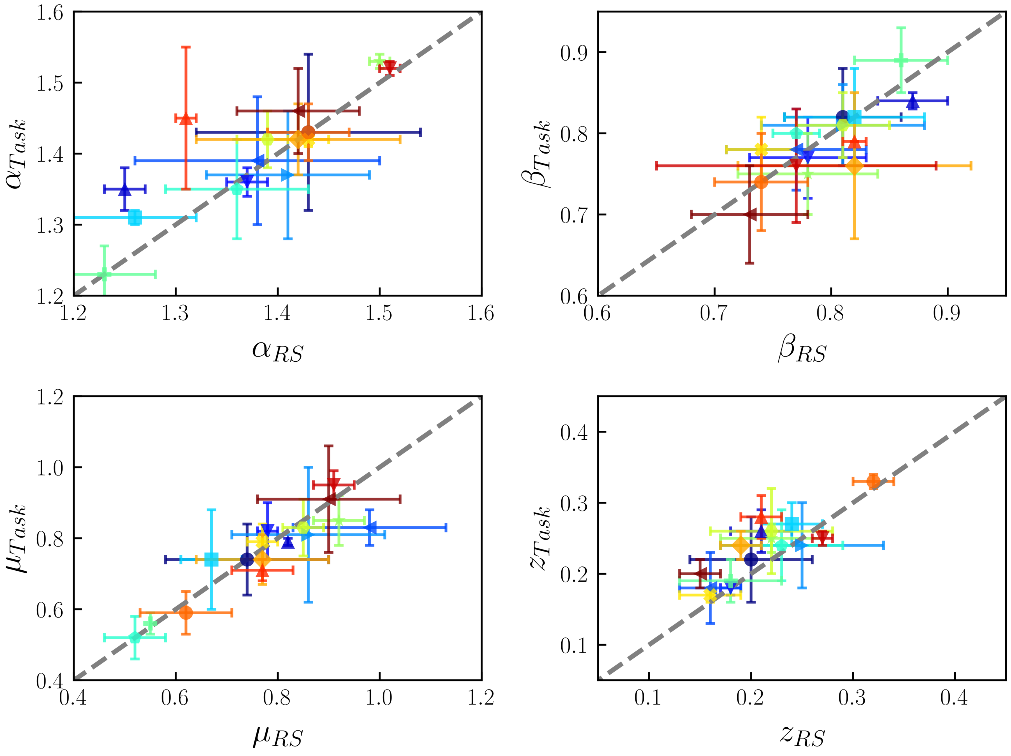


Figure 6: Scaling exponents for task-induced vs resting-state type of activity in 16 different regions of the mouse brain. Markers and colors for each area have been chosen as defined by Fig.1 in the main text. Error bars are defined as the standard deviation of the corresponding quantity across all experiments belonging to the same region.

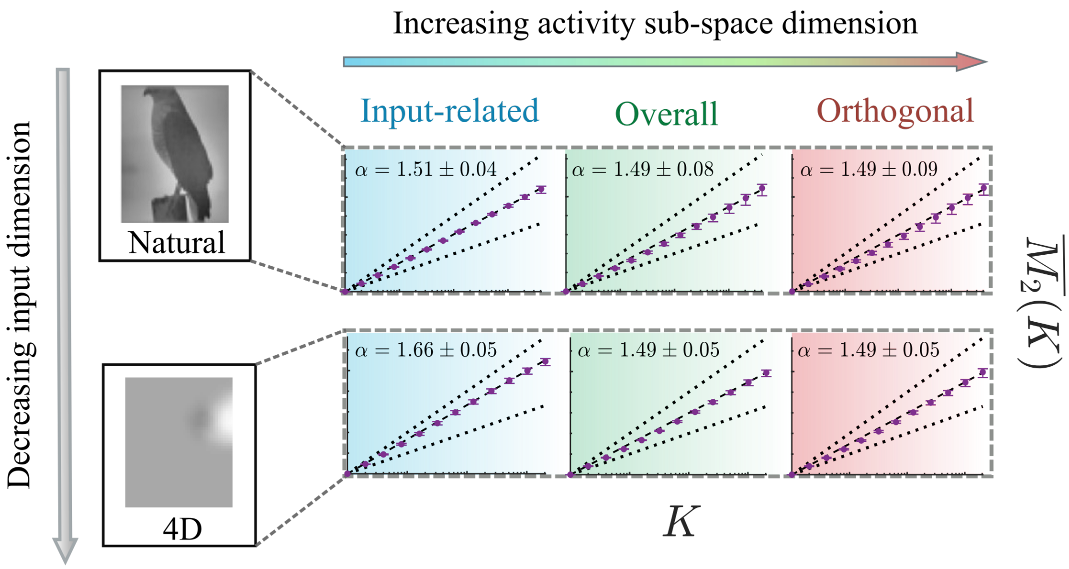


Figure 7: Diagram showing the two observed trends in the power-law exponent $\alpha$, which characterizes the scaling of the variance as a function of their rank. $\alpha$ increases with the complexity of the input when activity is projected into the task-encoding subspace (also called representation manifold). On the other hand, for a fixed type of input, the encoding activity shows a higher exponent (meaning higher correlations), while the background activity —which lies in a higher-dimensional subspace orthogonal to the representational manifold— and the overall/raw activity share the same exponents. Natural and low-dimensional images examples have been adapted from [2].

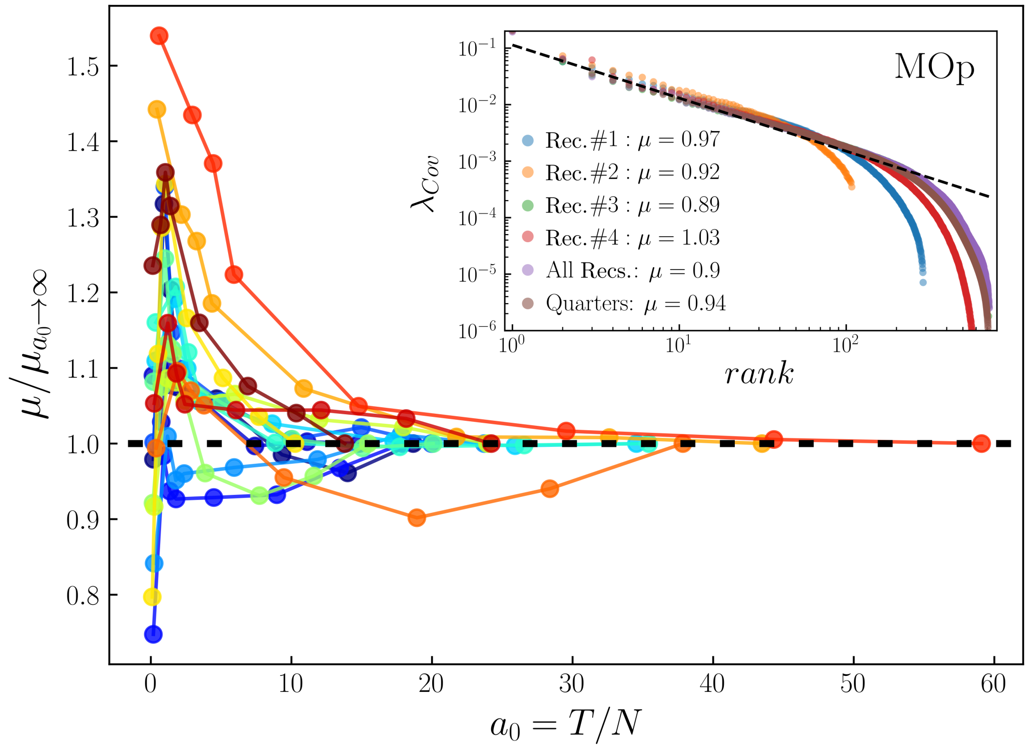


Figure 8: For each region, the scaling exponent for the covariance matrix eigenvalues at the last step of RG is plotted against the ratio samples-to-neurons in sub-sampled experiments, then re-scaled by its expected value when $a_0 \to \infty$ (estimated by using the full length of the time series). For each region, the experiment with the greatest number of recorded neurons was considered. **Inset:** Rank-ordered plot of eigenvalues in the MOp region using all the neurons while taking (i) each of the four recordings belonging to a experiment separately; (ii) merging them together; and (iii) averaging over quarter-splits of the data. Line fit is over quarters of the data.

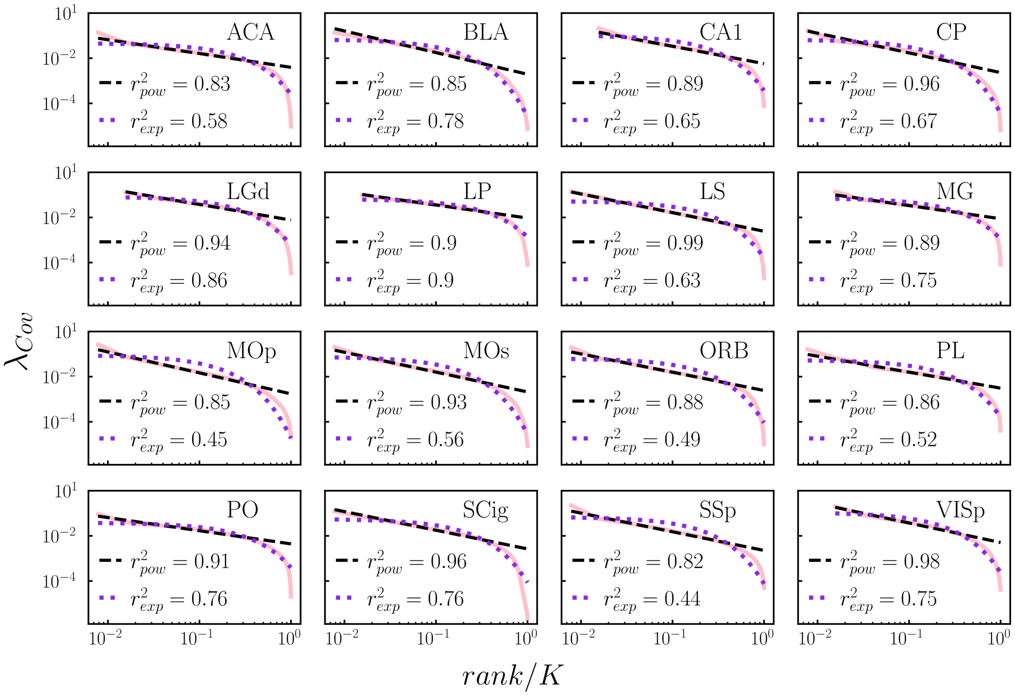


Figure 9: Best power-law (black dashed line) and exponential (purple dotted line) fits for the covariance matrix eigenvalues at the last step of RG (pink line), together with their corresponding R-Squared values. For each region, the experiment with the greatest number of recorded neurons was considered.

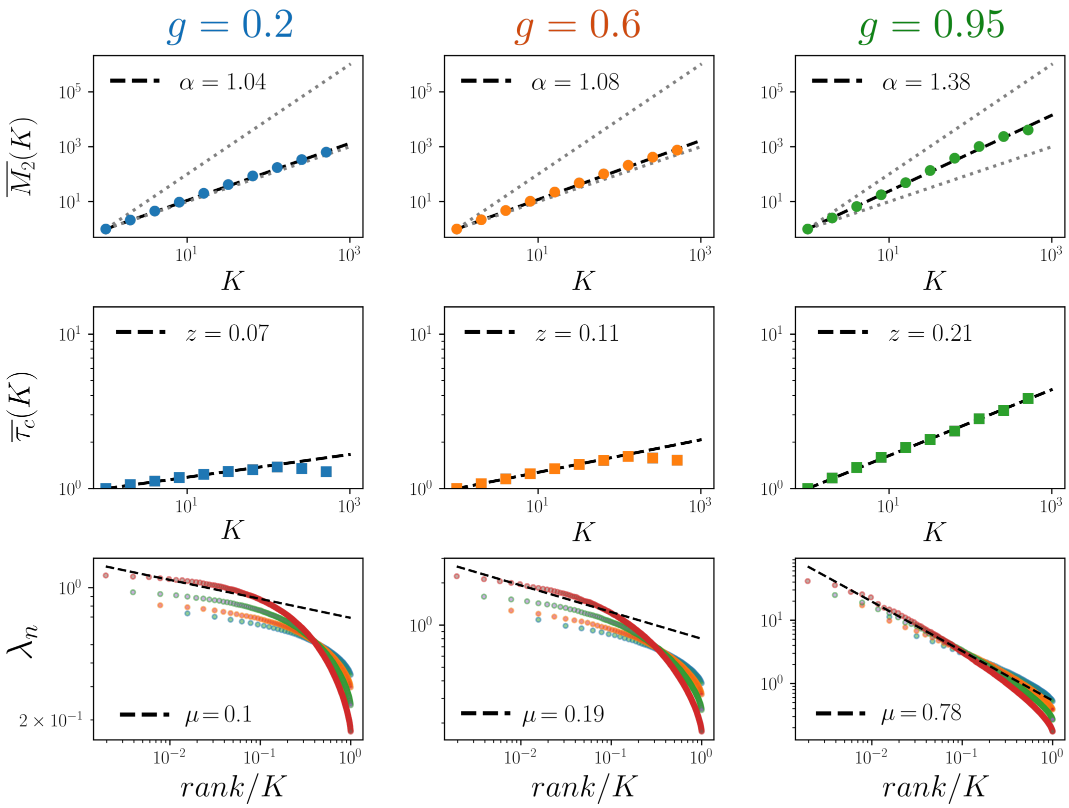


Figure 10: RG over randomly recurrent network of linear rate neurons for different values of the overall coupling strength $g$. Top: Variance of the non-normalized activity, Middle: scaling of the characteristic correlation time, Bottom: Covariance matrix spectrum for the resting-state activity in clusters of size $K$, as a function of the block-neuron size. Systems closer to the edge-of-instability ($g_c = 1$) show power-law scalings similar to the ones observed in the data. Neuron blocks in systems further away from such a regime behave like uncorrelated variables. In addition the rank-ordered spectrum is flatter and the scaling across RG steps is lost.

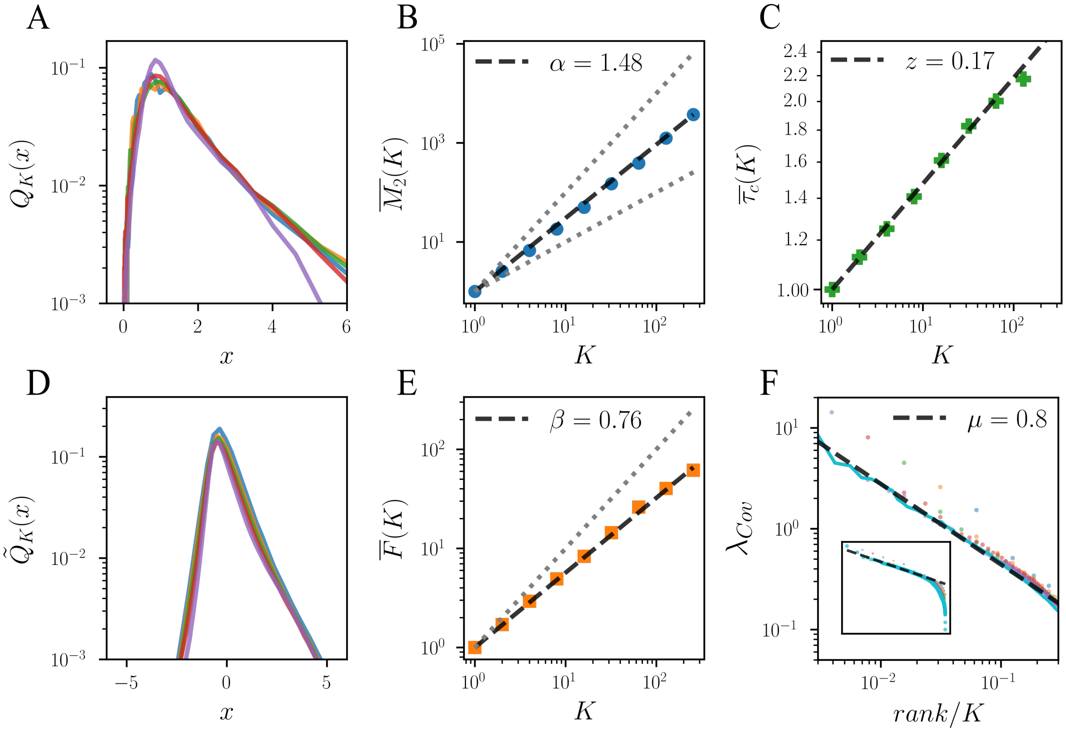


Figure 11: RG analysis over non-surrogated (control) data, for the region MOp. Plotted for comparison with surrogated reshuffled sets in Fig.12, 13 and 14. (A) Probability distribution for normalized non-zero activity in clusters of size $K = \{16, 32, 64, 128, 256\}$, corresponding to the yellow, blue, green, red and purple curves, respectively. (B) Scaling of activity variance inside clusters with cluster size. (C) Scaling of characteristic autocorrelation time with cluster size. (D) Probability distribution for normalized activity in momentum space, setting a cutoff on the first $K = \{16, 32, 64, 128, 256\}$ eigenmodes (computed as in [19], color code as in panel A). (E) Scaling of the "free-energy" with cluster size. (F) Inset: Covariance matrix spectrum with power law fit against the normalized rank. Main: closer view showing small-rank outliers.

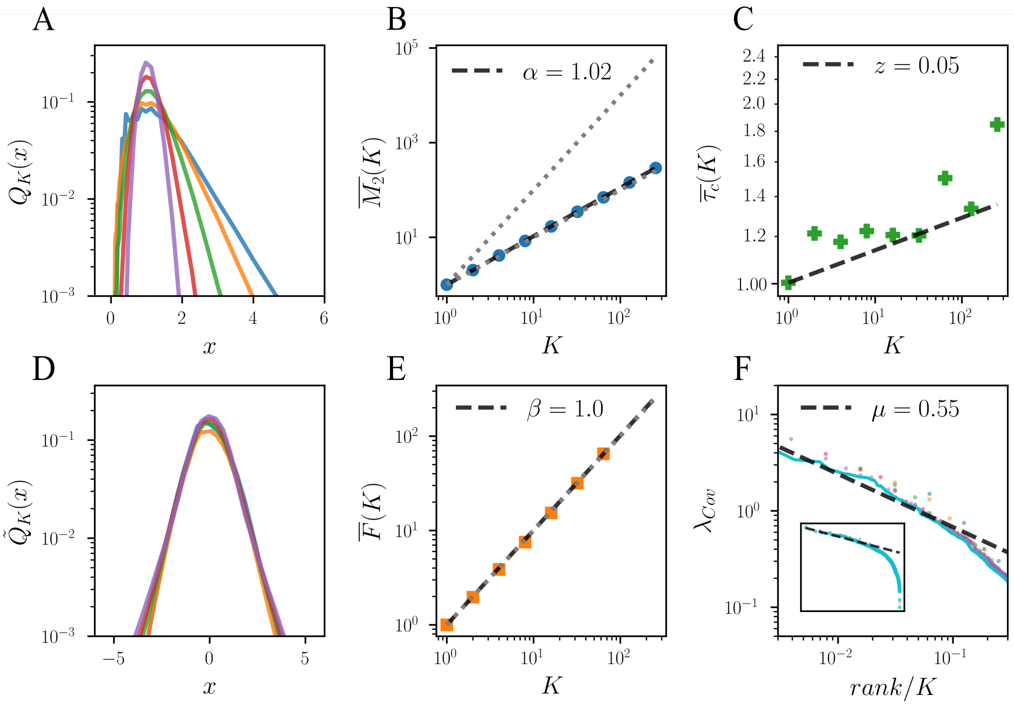


Figure 12: RG analysis over surrogated data created by by randomly shuffling all the spikes for each neuron. (A) Probability distribution for normalized non-zero activity in clusters of size $K = \{16, 32, 64, 128, 256\}$, corresponding to the yellow, blue, green, red and purple curves, respectively. (B) Scaling of activity variance inside clusters with cluster size. (C) Scaling of characteristic autocorrelation time with cluster size. (D) Probability distribution for normalized activity in momentum space, setting a cutoff on the first $K = \{16, 32, 64, 128, 256\}$ eigenmodes (computed as in [19], color code as in panel A) (E) Scaling of the "free-energy" with cluster size. (F) Inset: Covariance matrix spectrum with power law fit against the normalized rank. Main: closer view

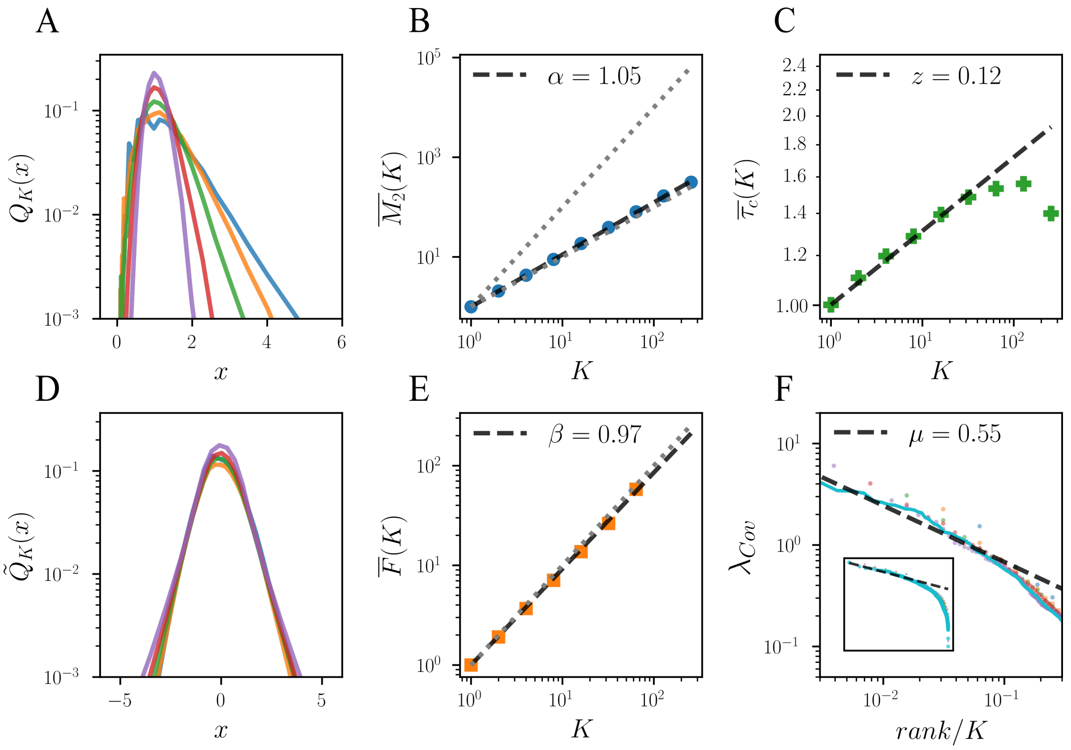


Figure 13: RG analysis over surrogated data created by shifting all the spikes of each neuron a random amount, so that the structure within the spike train is preserved. (A) Probability distribution for normalized non-zero activity in clusters of size $K = \{16, 32, 64, 128, 256\}$, corresponding to the yellow, blue, green, red and purple curves, respectively. (B) Scaling of activity variance inside clusters with cluster size. (C) Scaling of characteristic autocorrelation time with cluster size. (D) Probability distribution for normalized activity in momentum space, setting a cutoff on the first $K = \{16, 32, 64, 128, 256\}$ eigenmodes (computed as in [19], color code as in panel A) (E) Scaling of the "free-energy" with cluster size. (F) Inset: Covariance matrix spectrum with power law fit against the normalized rank. Main: closer view.

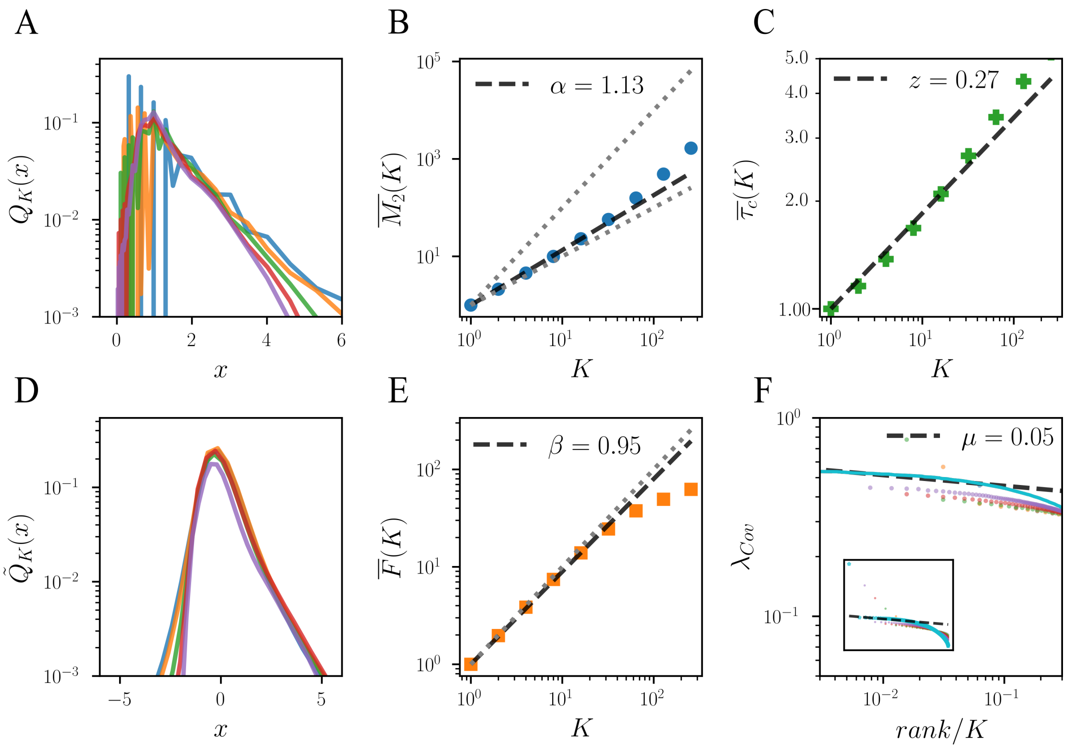


Figure 14: RG analysis over surrogated data created by randomizing the spikes across neurons but not time. (A) Probability distribution for normalized non-zero activity in clusters of size $K = \{16, 32, 64, 128, 256\}$, corresponding to the yellow, blue, green, red and purple curves, respectively. (B) Scaling of activity variance inside clusters with cluster size. (C) Scaling of characteristic autocorrelation time with cluster size. (D) Probability distribution for normalized activity in momentum space, setting a cutoff on the first $K = \{16, 32, 64, 128, 256\}$ eigenmodes (computed as in [19], color code as in panel A) (E) Scaling of the "free-energy" with cluster size. (F) Inset: Covariance matrix spectrum with (almost) flat power law fit against the normalized rank and strong first eigenvalue outlayer. Main: closer view

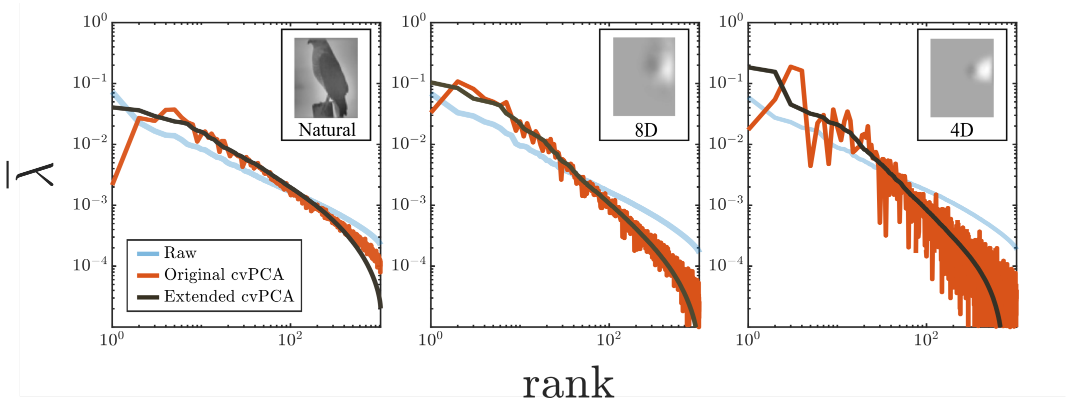


Figure 15: Distribution of pair-wise covariance eigenvalues (normalized as in [2] by their sum) measured from the activity of mice subject to three different types of stimuli: natural images (left panel), and their eight-dimensional (middle panel) and four-dimensional (right panel) projections. For each panel, the blue curve marks the spectrum measured on the overall, raw activity. Orange curves correspond to the spectrum of the input-related or encoding activity as measured by the cvPCA method proposed in [2]. On the other hand, black curves also represent the input-related covariance spectrum, but they are extracted directly from the proposed proxy for the input-related activity, as described in Section 3 of this SI.

Table 1: For each region we show: **(i)** the number $M$ of experiments considered; **(ii)** the average number of neurons across experiments; **(iii)** the average length of the recordings across experiments (once all intervals of spontaneous activity in an experiment have been merged together); **(iv)** the average lifetime and standard deviation across experiments of the correlations exponential decay, as given by the average autocorrelation function across all neurons in a region; **(v)** the average percentage of non-stationary neurons in a region; **(vi)** the distance to the edge of instability, as inferred in [15], with the standard deviation across experiments; **(vii)** same as in (vi), but using the method proposed in [14]; **(viii)** the error using the Cramer-von Mises statistic between the sampled eigenvalue distribution and the theoretical distribution proposed in [14] for a model of linear rate neurons with recurrent connectivity (RC); **(ix)** same as in (viii), but comparing the empirical distribution to a best-fitting Marchenko-Pastur (MP) distribution; **(x)** estimated distance to the edge of instability using a common extrapolated size of $N = 10^4$ neurons, with the error given as the standard deviation across experiments. For a given region, each experiment corresponds to a different mouse, and only mice having more than $N = 128$ recorded were selected.

| Full name | Abbrev. | $M$ | $N\left(\times 10^2\right)$ | $T\left(\times 10^2 s\right)$ | $\tau_{Corr}(ms)$ | % N.S. | $\lambda_{max}$ | $g$ | $L^2_{RC}$ | $L^2_{MP}$ | $\lambda_{max}^{N=1e4}$ |
|---|---|---|---|---|---|---|---|---|---|---|---|
| Anterior cingulate area | ACA | 3 | $1.8 \pm 0.8$ | $7.5 \pm 3.2$ | $0.10 \pm 0.02$ | $1.2 \pm 0.5$ | $0.70 \pm 0.09$ | $0.74 \pm 0.09$ | $0.07 \pm 0.02$ | $0.11 \pm 0.04$ | $0.965 \pm 0.013$ |
| Basolateral amygdalar nucleus | BLA | 2 | $2.7 \pm 0.0$ | $8.1 \pm 0.5$ | $0.09 \pm 0.01$ | $1.3 \pm 0.6$ | $0.81 \pm 0.00$ | $0.86 \pm 0.04$ | $0.07 \pm 0.02$ | $0.17 \pm 0.02$ | $0.969 \pm 0.000$ |
| Cornu ammonis | CA1 | 2 | $1.9 \pm 0.2$ | $8.8 \pm 2.9$ | $0.07 \pm 0.01$ | $7.7 \pm 0.3$ | $0.85 \pm 0.04$ | $0.83 \pm 0.05$ | $0.06 \pm 0.01$ | $0.15 \pm 0.02$ | $0.980 \pm 0.004$ |
| Caudoputamen | CP | 4 | $4.1 \pm 1.0$ | $11.0 \pm 1.2$ | $0.09 \pm 0.01$ | $2.6 \pm 1.4$ | $0.91 \pm 0.02$ | $0.93 \pm 0.01$ | $0.09 \pm 0.02$ | $0.23 \pm 0.01$ | $0.983 \pm 0.002$ |
| Lateral septal nucleus | LS | 3 | $2.6 \pm 0.5$ | $6.4 \pm 2.9$ | $0.05 \pm 0.03$ | $2.1 \pm 0.5$ | $0.90 \pm 0.02$ | $0.83 \pm 0.02$ | $0.05 \pm 0.00$ | $0.15 \pm 0.00$ | $0.972 \pm 0.007$ |
| Dorsal part of the lateral geniculate complex | LGd | 5 | $1.6 \pm 0.1$ | $8.3 \pm 1.0$ | $0.08 \pm 0.01$ | $1.6 \pm 1.6$ | $0.77 \pm 0.04$ | $0.82 \pm 0.06$ | $0.09 \pm 0.02$ | $0.12 \pm 0.03$ | $0.962 \pm 0.009$ |
| Lateral posterior nucleus of the thalamus | LP | 2 | $1.8 \pm 0.1$ | $9.4 \pm 0.7$ | $0.08 \pm 0.01$ | $0.3 \pm 0.3$ | $0.72 \pm 0.06$ | $0.66 \pm 0.06$ | $0.07 \pm 0.04$ | $0.06 \pm 0.00$ | $0.983 \pm 0.002$ |
| Medial geniculate complex of the thalamus | MG | 2 | $2.2 \pm 0.2$ | $10.0 \pm 2.4$ | $0.09 \pm 0.01$ | $3.4 \pm 3.4$ | $0.74 \pm 0.07$ | $0.73 \pm 0.00$ | $0.09 \pm 0.02$ | $0.08 \pm 0.01$ | $0.962 \pm 0.009$ |
| Primary motor area | MOp | 3 | $4.2 \pm 2.2$ | $11.5 \pm 0.9$ | $0.12 \pm 0.02$ | $1.3 \pm 0.5$ | $0.93 \pm 0.01$ | $0.91 \pm 0.02$ | $0.09 \pm 0.01$ | $0.27 \pm 0.02$ | $0.987 \pm 0.002$ |
| Secondary motor area | MOs | 5 | $2.5 \pm 0.7$ | $8.5 \pm 3.2$ | $0.10 \pm 0.01$ | $4.3 \pm 1.8$ | $0.84 \pm 0.02$ | $0.84 \pm 0.03$ | $0.06 \pm 0.01$ | $0.16 \pm 0.02$ | $0.975 \pm 0.000$ |
| Orbital area | ORB | 3 | $2.7 \pm 0.3$ | $9.5 \pm 0.8$ | $0.09 \pm 0.02$ | $1.8 \pm 0.7$ | $0.87 \pm 0.02$ | $0.80 \pm 0.02$ | $0.05 \pm 0.01$ | $0.13 \pm 0.01$ | $0.980 \pm 0.003$ |
| Prelimbic area | PL | 4 | $2.4 \pm 0.5$ | $8.7 \pm 1.6$ | $0.12 \pm 0.04$ | $3.9 \pm 2.3$ | $0.83 \pm 0.05$ | $0.80 \pm 0.08$ | $0.04 \pm 0.01$ | $0.14 \pm 0.05$ | $0.974 \pm 0.010$ |
| Posterior complex of the thalamus | PO | 3 | $2.8 \pm 1.1$ | $10.1 \pm 1.0$ | $0.10 \pm 0.03$ | $0.9 \pm 0.6$ | $0.89 \pm 0.02$ | $0.78 \pm 0.05$ | $0.09 \pm 0.02$ | $0.12 \pm 0.03$ | $0.983 \pm 0.005$ |
| Superior culliculus, intermediate gray layer | SCig | 2 | $2.8 \pm 0.9$ | $8.4 \pm 0.4$ | $0.11 \pm 0.00$ | $3.6 \pm 2.0$ | $0.91 \pm 0.01$ | $0.86 \pm 0.05$ | $0.08 \pm 0.02$ | $0.19 \pm 0.04$ | $0.986 \pm 0.001$ |
| Primary somatosensory area | SSp | 2 | $3.3 \pm 0.0$ | $11.6 \pm 1.0$ | $0.12 \pm 0.01$ | $1.7 \pm 0.7$ | $0.93 \pm 0.01$ | $0.91 \pm 0.02$ | $0.08 \pm 0.01$ | $0.20 \pm 0.02$ | $0.988 \pm 0.002$ |
| Primary visual area | VISp | 4 | $1.9 \pm 0.3$ | $9.5 \pm 2.1$ | $0.08 \pm 0.01$ | $4.0 + 2.5$ | $0.84 \pm 0.03$ | $0.85 \pm 0.02$ | $0.07 \pm 0.00$ | $0.16 \pm 0.02$ | $0.978 \pm 0.004$ |

Table 2: For each region we show the average bin size $\Delta t$ used in the RG analysis (with standard deviation across experiments) as given by the geometric mean of the ISIS distribution. Moreover, for each region and measured exponents we collect: **(i)** the average value across experiments; **(ii)** the mean-absolute-error, computed as the average across experiments of the experiment-specific errors measured over split-quarters of data; **(iii)** the standard deviation across experiments; **(iv)** the R-squared value of the best powerlaw fit for the experiment with a greater number of recorded neurons. For the exponent of the rank-ordered covariance matrix eigenvalues, we also provide the R-squared value of the best exponential fit, as well as the likelihood ratios and p-values for the significance of each test comparing the powerlaw fit of the associated eigenvalue density with corresponding exponential and lognormal distributions (statistically significant p-values are highlighted in boldface). Positive ratios indicate that the data is best fitted by a powerlaw distribution. For a given region, each experiment corresponds to a different mouse, and only mice having more than $N = 128$ recorded neurons were selected.

| Abbrev. | $\Delta t(s)$ | $\langle \boldsymbol{\alpha} \rangle$ | MAE | $\sigma$ | $r^2$ | $\langle \boldsymbol{\beta} \rangle$ | MAE | $\sigma$ | $r^2$ | $\langle \mathbf{z} \rangle$ | MAE | $\sigma$ | $r^2$ | $\langle \boldsymbol{\mu} \rangle$ | MAE | $\sigma$ | $r^2_{\text{pow}}$ | $r^2_{\text{exp}}$ | Exponential | | Lognormal | |
|---|---|---|---|---|---|---|---|---|---|---|---|---|---|---|---|---|---|---|---|---|---|---|
| | | | | | | | | | | | | | | | | | | | LR | $p$ | LR | $p$ |
| ACA | $0.16 \pm 0.07$ | 1.43 | 0.02 | 0.11 | 1.00 | 0.81 | 0.03 | 0.05 | 1.00 | 0.2 | 0.02 | 0.06 | 0.98 | 0.74 | 0.05 | 0.16 | 0.83 | 0.58 | 90.5 | **0.00** | -0.1 | 0.22 |
| BLA | $0.13 \pm 0.01$ | 1.25 | 0.03 | 0.02 | 1.00 | 0.87 | 0.04 | 0.03 | 1.00 | 0.21 | 0.03 | 0.03 | 1.00 | 0.82 | 0.03 | 0.00 | 0.85 | 0.78 | 9.4 | 0.12 | -2.9 | 0.09 |
| CA1 | $0.13 \pm 0.04$ | 1.37 | 0.03 | 0.02 | 1.00 | 0.78 | 0.04 | 0.05 | 1.00 | 0.18 | 0.03 | 0.01 | 0.99 | 0.78 | 0.08 | 0.02 | 0.89 | 0.65 | 55.4 | **0.00** | -0.3 | 0.59 |
| CP | $0.08 \pm 0.02$ | 1.38 | 0.02 | 0.12 | 1.00 | 0.77 | 0.05 | 0.06 | 0.98 | 0.16 | 0.02 | 0.03 | 0.99 | 0.98 | 0.1 | 0.15 | 0.96 | 0.67 | 43.3 | **0.00** | -2.5 | 0.17 |
| LS | $0.17 \pm 0.13$ | 1.41 | 0.05 | 0.08 | 1.00 | 0.81 | 0.04 | 0.07 | 1.00 | 0.25 | 0.05 | 0.08 | 0.99 | 0.86 | 0.08 | 0.15 | 0.94 | 0.86 | 8.4 | **0.05** | -0.9 | 0.32 |
| LGd | $0.07 \pm 0.03$ | 1.26 | 0.03 | 0.06 | 1.00 | 0.82 | 0.04 | 0.06 | 1.00 | 0.24 | 0.06 | 0.03 | 1.00 | 0.67 | 0.03 | 0.06 | 0.9 | 0.9 | 0.5 | 0.86 | -1.5 | 0.21 |
| LP | $0.08 \pm 0.02$ | 1.36 | 0.03 | 0.07 | 1.00 | 0.77 | 0.02 | 0.02 | 1.00 | 0.23 | 0.05 | 0.06 | 0.99 | 0.52 | 0.06 | 0.06 | 0.99 | 0.63 | 152.3 | **0.00** | -3.5 | 0.08 |
| MG | $0.06 \pm 0.01$ | 1.23 | 0.02 | 0.05 | 1.00 | 0.86 | 0.02 | 0.04 | 1.00 | 0.18 | 0.04 | 0.05 | 0.99 | 0.55 | 0.05 | 0.00 | 0.89 | 0.75 | 24.3 | **0.00** | -0.0 | 0.90 |
| MOp | $0.06 \pm 0.01$ | 1.5 | 0.03 | 0.01 | 1.00 | 0.78 | 0.03 | 0.06 | 0.99 | 0.22 | 0.02 | 0.05 | 1.00 | 0.92 | 0.05 | 0.05 | 0.85 | 0.45 | 138.8 | **0.00** | -0.2 | 0.66 |
| MOs | $0.16 \pm 0.03$ | 1.39 | 0.04 | 0.03 | 1.00 | 0.81 | 0.04 | 0.04 | 1.00 | 0.22 | 0.03 | 0.06 | 0.99 | 0.85 | 0.07 | 0.04 | 0.93 | 0.56 | 85.1 | **0.00** | -0.0 | 0.87 |
| ORB | $0.12 \pm 0.02$ | 1.43 | 0.02 | 0.02 | 1.00 | 0.74 | 0.02 | 0.03 | 0.99 | 0.16 | 0.02 | 0.03 | 0.99 | 0.77 | 0.03 | 0.03 | 0.88 | 0.49 | 146.5 | **0.00** | -0.6 | 0.50 |
| PL | $0.13 \pm 0.08$ | 1.42 | 0.04 | 0.10 | 1.00 | 0.82 | 0.04 | 0.1 | 0.99 | 0.19 | 0.03 | 0.02 | 0.99 | 0.77 | 0.05 | 0.13 | 0.86 | 0.52 | 180.9 | **0.00** | -0.6 | 0.48 |
| PO | $0.07 \pm 0.02$ | 1.43 | 0.02 | 0.04 | 1.00 | 0.74 | 0.03 | 0.04 | 1.00 | 0.32 | 0.01 | 0.02 | 0.99 | 0.62 | 0.03 | 0.09 | 0.91 | 0.76 | 22.8 | **0.00** | 0.0 | 0.85 |
| SCig | $0.07 \pm 0.00$ | 1.31 | 0.04 | 0.01 | 1.00 | 0.82 | 0.04 | 0.01 | 0.98 | 0.21 | 0.04 | 0.02 | 1.00 | 0.77 | 0.08 | 0.06 | 0.96 | 0.76 | 26.6 | **0.00** | -2.8 | 0.11 |
| SSp | $0.06 \pm 0.01$ | 1.51 | 0.04 | 0.01 | 1.00 | 0.77 | 0.02 | 0.12 | 0.99 | 0.27 | 0.02 | 0.01 | 1.00 | 0.91 | 0.03 | 0.04 | 0.82 | 0.44 | 121.4 | **0.00** | -0.5 | 0.54 |
| VISp | $0.12 \pm 0.04$ | 1.42 | 0.02 | 0.06 | 1.00 | 0.73 | 0.02 | 0.05 | 1.00 | 0.15 | 0.02 | 0.02 | 0.98 | 0.90 | 0.04 | 0.14 | 0.98 | 0.75 | 22.2 | **0.00** | -1.2 | 0.26 |

[14] Yu Hu and Haim Sompolinsky. The spectrum of covariance matrices of randomly connected recurrent neuronal networks. *bioRxiv*, 2020.

[15] David Dahmen, Sonja Grün, Markus Diesmann, and Moritz Helias. Second type of criticality in the brain uncovers rich multiple-neuron dynamics. *Proceedings of the National Academy of Sciences*, 116(26):13051–13060, June 2019.

[16] Weihao Kong and Gregory Valiant. Spectrum estimation from samples. *The Annals of Statistics*, 45(5):2218–2247, October 2017. Publisher: Institute of Mathematical Statistics.

[17] Giordano De Marzo, Andrea Gabrielli, Andrea Zaccaria, and Luciano Pietronero. Dynamical approach to zipf's law. *Physical Review Research*, 3(1):013084, 2021.

[18] Wentian Li. Zipf's law everywhere. *Glottometrics*, 5:14–21, 2002.

[19] Leenoy Meshulam, Jeffrey L Gauthier, Carlos D Brody, David W Tank, and William Bialek. Coarse–graining and hints of scaling in a population of 1000+ neurons. *arXiv preprint arXiv:1812.11904*, 2018.
